# CYB5R3 Controls Sex-Specific Stress Erythropoiesis via Heme-Biosynthesis

**DOI:** 10.64898/2025.12.04.692104

**Authors:** Fabliha A Chowdhury, Malini Sharma, Katherine C Wood, Sarah A Saad, Shuai Yuan, Megan P Miller, Scott A Hahn, Mate Katona, Stefanie N Taiclet, Sonia R Salvatore, Francisco J Schopfer, Adam C Straub

## Abstract

Cytochrome b5 reductase 3 (CYB5R3) or met-hemoglobin reductase is an oxidoreductase that maintains hemoprotein and cellular redox balance, yet its contribution to erythropoiesis under stress conditions remains unclear. Motivated by prior observations that the hypomorphic CYB5R3 T117S blunts hydroxyurea-induced fetal hemoglobin responses in patients with sickle cell disease, we tested whether CYB5R3 contributes to the regulation of erythropoiesis. Hematopoietic lineage-specific *CYB5R3* knockout mice exhibited markedly impaired erythropoietic induction in response to chronic hypoxia compared to controls, with males showing a more pronounced deficit, and splenectomy further exacerbating this impairment. Genetic deletion of *CYB5R3* in human CD34⁺ progenitors reduced globin expression and disrupted terminal erythroid differentiation. Meanwhile, *CYB5R3* knockdown in K562 cells produced a heme-deficient state whereby only exogenous heme but not hydroxyurea, iron, or upstream precursors restored globin synthesis. Transcriptomic profiling revealed coordinated downregulation of erythroid transcription factors and multiple enzymes in the heme biosynthetic pathway, which was reversed with heme treatment. Together, these results reveal an unexpected function for CYB5R3 beyond met-hemoglobin reduction, positioning it as a central metabolic regulator of sex-specific stress erythropoiesis and unveiling a heme-restricted vulnerability that may augment disease severity in anemia, hemoglobinopathies, and individuals carrying CYB5R3 loss-of-function variants.

**Key points:**

- CYB5R3 is required for effective stress erythropoietic induction, with a more pronounced impact in males.
- Erythroid-specific CYB5R3 deficiency creates a heme-limited state, impairing erythroblast differentiation and maturation.

## Introduction

Anemia, defined as a deficiency of mature and healthy erythrocytes, is strongly associated with poor health outcomes, increased health expenses, and elevated morbidity and mortality rates worldwide^1^. With a global burden of 1.92 billion in 2021, anemia primarily affects minors, women of reproductive age, and the elderly^1^. Although significant advances have been made in distinguishing classical and pathological erythropoiesis, there remains a substantial gap in developing appropriate anemia-targeted interventions to alleviate disease burden^1,2^. Compelling evidence indicates that oxidative stress resulting from altered redox signaling significantly impairs erythrocyte formation, function, and survival^3,4^. While the mechanism remains incompletely understood, impaired redox balance and anemia are found to be extensively enhanced in various systemic phenomena, including inflammation, diabetes, malignancy, and chronic heart, liver, and kidney diseases^3^.

The human body generates approximately 2 million erythrocytes every second through a process known as erythropoiesis, which occurs in the bone marrow at steady-state but can extend to the spleen during stress^5^. The principal content of a mature erythrocyte is hemoglobin, a tetramer composed of four globin chains, each containing a heme group^6,7,8^. Functionally, heme-containing hemoglobin in the erythrocytes binds and carries oxygen in the circulation in a reduced ferrous state^9^. This heme iron can oxidize to its ferric state (met-hemoglobin) under physiological or pathological conditions, becoming incapable of binding oxygen^9,10^. Cytochrome b5 reductase 3 (CYB5R3), also known as met-hemoglobin reductase, reduces heme back to its active ferrous state^11–12^. It is well-established that CYB5R3 plays a critical role in preserving the redox equilibrium in physiology and disease. CYB5R3 is usually found in the cytoplasm of mature erythrocytes, whereas a membrane-bound form exists in all other cell types^11,13,14^. Heme reduction by CYB5R3 extends beyond hemoglobin to soluble guanylate cyclase (sGC), where it reduces ferric to ferrous heme, thereby modulating nitric oxide (NO)-sGC-cyclic GMP (cGMP) signaling^13,15^. The NO-sGC-cGMP pathway plays an essential role in various physiological processes, including erythropoiesis, linking it to CYB5R3^16,17^.

Approximately 40 genetic variants of CYB5R3 have been reported^12,18^. Among them, the CYB5R3 T117S variant is highly relevant to humans as it has ∼50% decreased reductase activity and a high-frequency expression (0.23 minor allele) in persons of African ancestry^13,19,20^. Interestingly, the same population is predominantly afflicted with sickle cell disease (SCD).^21^ Our recent study analyzing the Walk-PHaSST clinical trial data demonstrated that, compared to wild-type, CYB5R3 T117S reduces fetal hemoglobin (HbF) induction by over 30% in SCD patients undergoing hydroxyurea (HU) treatment.^19^ Additionally, CYB5R3 deletion in primary CD34+ hematopoietic stem cells (HSCs) impaired HU-induced HbF production and erythroid maturation *in vitro*.^19^ While emerging evidence suggests a potential new role for CYB5R3 in hemoglobin synthesis and erythropoiesis, the underlying mechanistic underpinnings remain unclear. Hence, based on our aforementioned findings and published data, we hypothesized that CYB5R3 is essential for erythropoiesis.

## Methodology

### Animal experiment

Erythropoiesis is a multistep process that initiates from HSCs **(Supp. Figure 1A)**. To check the role of CYB5R3 in erythropoiesis *in vivo*, female *CYB5R3 f/+* mice were crossed with male *Vav1Cre* mice (Jackson Laboratory, 035670), both on the C57BL/6J background **(Supp. Figure 1B)** ^22^. The *Vav1Cre* recombinase is expressed in HSCs and all nucleated blood cells under the regulation of the human Vav1 oncogene regulatory elements, which can be targeted for deletion of the floxed gene in the entire hematopoietic compartment.^23,24^ Heterozygous *CYB5R3f/+;Vav1Cre+/-* littermates were crossed to generate heterozygous *Cre*-positive, *CYB5R3* homozygous knockout (*CYB5R3f/f;Vav1Cre+/-, CYB5R3* KO) mice. Littermates, *CYB5R3 f/f* mice without *Vav1Cre,* were used as wild-type (WT) controls. The deletion of the *CYB5R3* gene was confirmed using DNA genotyping and Western Blot **(Supp. Figure 1C)**. 12-16 weeks old adult mice, both males and females, were used for baseline, hypoxia, and splenectomy studies.

**Figure 1.**
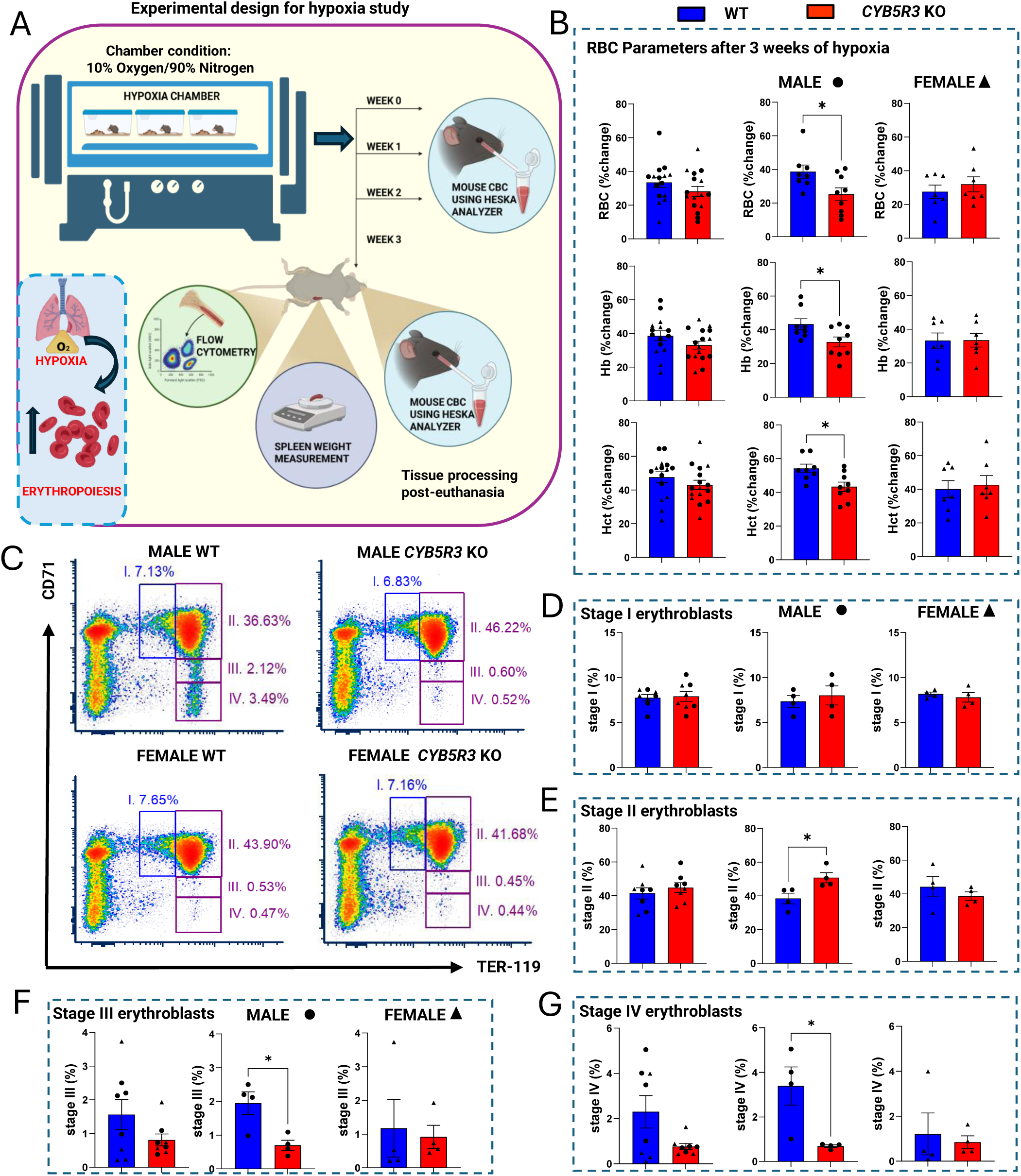
Absence of hematopoietic lineage-specific CYB5R3 lowers the erythropoietic induction by hypoxia in male mice through an erythroblast differentiation defect. A. Experimental design for hypoxia study. B. Percentage change in erythrocyte (RBC) parameters after 3 weeks of hypoxia, combined, and males and females separately. C. Flow cytometry plots for erythroblast stages in the bone marrow after 3 weeks of hypoxia. D-G. Quantification of different stages of erythroblasts in the bone marrow, combined, and males and females separately. Flow cytometry was performed using a BD Biosciences LSR II. Data were analyzed using FCS Express 7 Research Edition (De Novo Software). GraphPad Prism version 10 was used for statistical analyses. Data are stated as mean ± standard error of the mean unless indicated otherwise. Parametric and non-parametric data with 1 variable were analyzed using Student’s t-test and Mann-Whitney test, respectively. F-test was used to compare variances. Grouped data with 2 variables were analyzed using 2-way ANOVA. * indicates p<0.05 and was considered significant. Data are stated as mean ± standard error of the mean unless indicated otherwise. CYB5R3: Cytochrome B5 reductase 3; WT: Wild-type; KO: Knockout; RBC: red blood cells, erythrocytes; Hb: hemoglobin; Hct: hematocrit

### CD34+ cell experiment

Healthy human CD34+ HSCs were acquired from Fred Hutchinson Cancer Research Center. The cells were cultured in 3 different phases as mentioned previously.^19^ On day 3 of cell culture, CD34+ HSCs underwent electroporation to generate *CYB5R3* KO cells or non-targeting (NT) control cells, using Cas9 proteins and guide RNAs. Cells were collected on day 11 for gene expression analysis, on day 15 for performing Western Blot, and on day 18 for flow cytometry.

### K562 cell experiment

K562 cells, a human erythroleukemic cell line, were cultured in RPMI 1640 supplemented with 10% fetal bovine serum (FBS) and 1% Penicillin/Streptomycin^25,26^. CYB5R3 was deleted in K562 cells using the lentiviral transduction method to generate CYB5R3 knockdown (*CYB5R3* KD) and NT control cells. Afterward, the cells were treated with different agents, followed by 6 hours to check globin mRNA induction and 48 hours to check globin protein induction. The treatment agents and dosages used are as follows: 50 µM HU, 10 µM hemin, 25 µM ferric chloride (FeCl_3_), and 50 µM 5-aminolevulinic acid (ALA). DMSO was used as the vehicle.

### Statistics

We used GraphPad Prism version 10 for statistical analyses. Parametric data were analyzed using Student’s t-test and non-parametric data using Mann-Whitney test, for 1 variable analysis. F-test was used to compare variances. 2-way ANOVA was used to analyze grouped data with 2 variables. A value of p<0.05 was considered significant.

## Results

### CYB5R3 in the hematopoietic compartment is dispensable for steady-state

***erythropoiesis.*** We discovered that *CYB5R3* KO adult mice had a darker color of blood, consistent with met-hemoglobinemia, relative to controls **(Supp. Figure 1C).** Hematological parameters in peripheral blood and bone marrow were measured **(Supp. Figure 2A)**. We did not observe any significant differences in the erythrocyte counts (RBC), hemoglobin level (Hb), or hematocrit (Hct) **(Supp. Figure 2B)** between the two groups, regardless of sex. Although the platelet counts were slightly higher in the female *CYB5R3* KOs relative to female WTs **(Supp. Figure 3A)**, the white blood cell (WBC) counts were similar **(Supp. Figure 3B)**. The post-mortem spleen size did not differ between groups, regardless of sex **(Supp. Figure 3C).** Different stages of erythroblast populations were measured from the bone marrow cells using flow cytometry, with only Stage IV erythroblasts showing a significant increase in the female KO mice **(Supp. Figures 2C-G)**. The HSCs and common myeloid progenitors (CMPs) in the bone marrow showed no significant differences between groups, regardless of sex **(Supp. Figures 3D-G)**. The megakaryocyte erythroid progenitors (MEPs) were comparatively higher, and granulocyte monocyte progenitors (GMPs) were lower in the male *CYB5R3* KOs **(Supp. Figures 3F-I)**; however, this did not really affect the hematological parameters compared to male WTs. Together, the data showed that *CYB5R3* KO mice exhibited normal erythropoiesis at steady-state.

**Figure 2.**
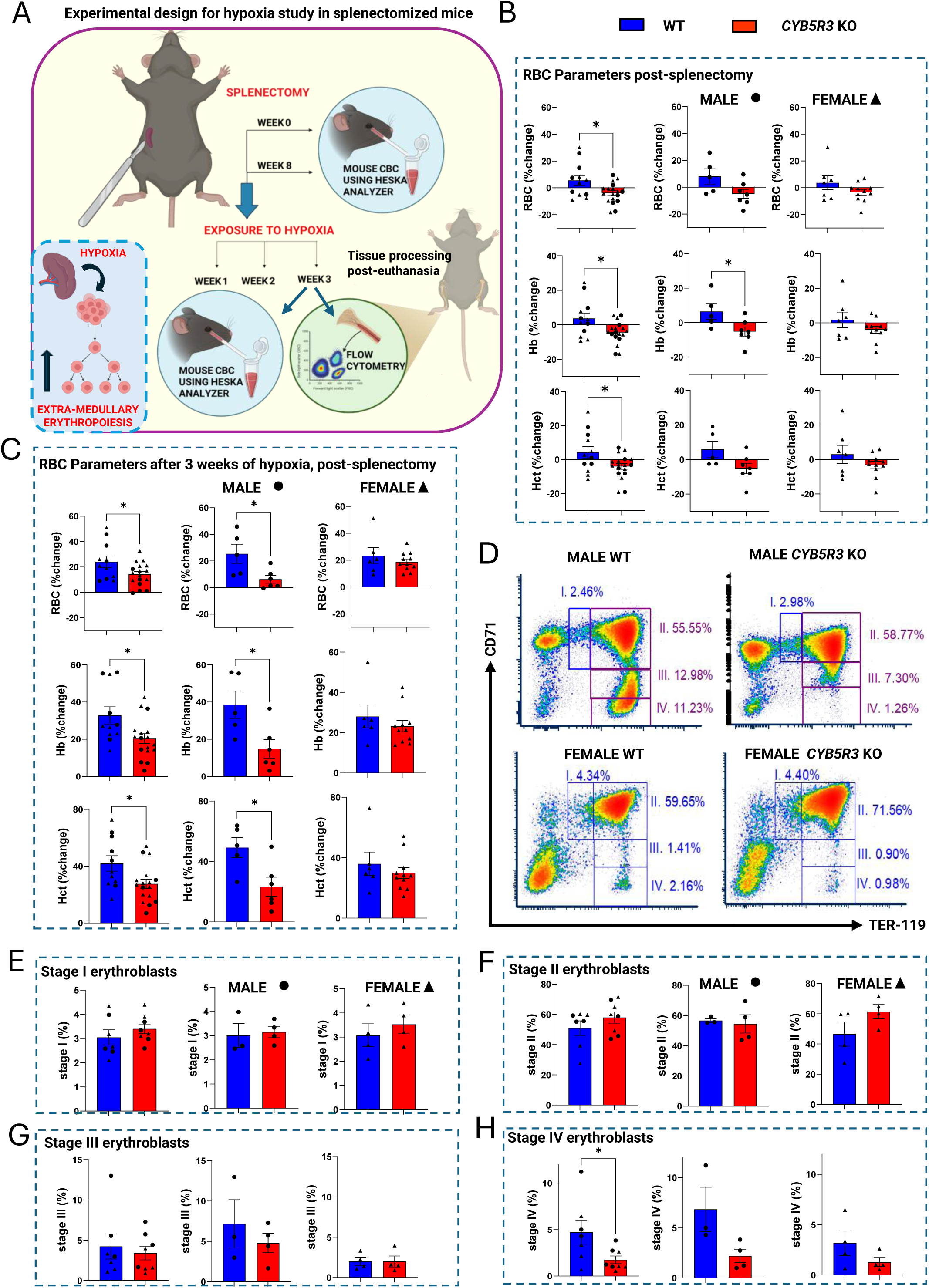
Splenectomy induces a significant drop in erythropoiesis in *CYB5R3* KO mice, which further impacts erythropoietic induction by hypoxia, specifically in male mice. A. Experimental design for hypoxia study in splenectomized mice. **B.** Percentage change in erythrocyte (RBC) parameters post-splenectomy, combined, and males and females separately. **C.** Percentage change in erythrocyte (RBC) parameters after 3 weeks of hypoxia, post-splenectomy, combined, and males and females separately. **D.** Flow cytometry plots for erythroblast stages in the bone marrow after 3 weeks of hypoxia, post-splenectomy. **E-H.** Quantification of different stages of erythroblasts in the bone marrow, combined, and males and females separately. Flow cytometry was performed using a BD Biosciences LSR II. Data were analyzed using FCS Express 7 Research Edition (De Novo Software). GraphPad Prism version 10 was used for statistical analyses. Data are stated as mean ± standard error of the mean unless indicated otherwise. Parametric and non-parametric data with 1 variable were analyzed using Student’s t-test and Mann-Whitney test, respectively. F-test was used to compare variances. Grouped data with 2 variables were analyzed using 2-way ANOVA. * indicates p<0.05 and was considered significant. CYB5R3: Cytochrome B5 reductase 3; WT: Wild-type; KO: Knockout; RBC: red blood cells, erythrocytes; Hb: hemoglobin; Hct: hematocrit

**Figure 3.**
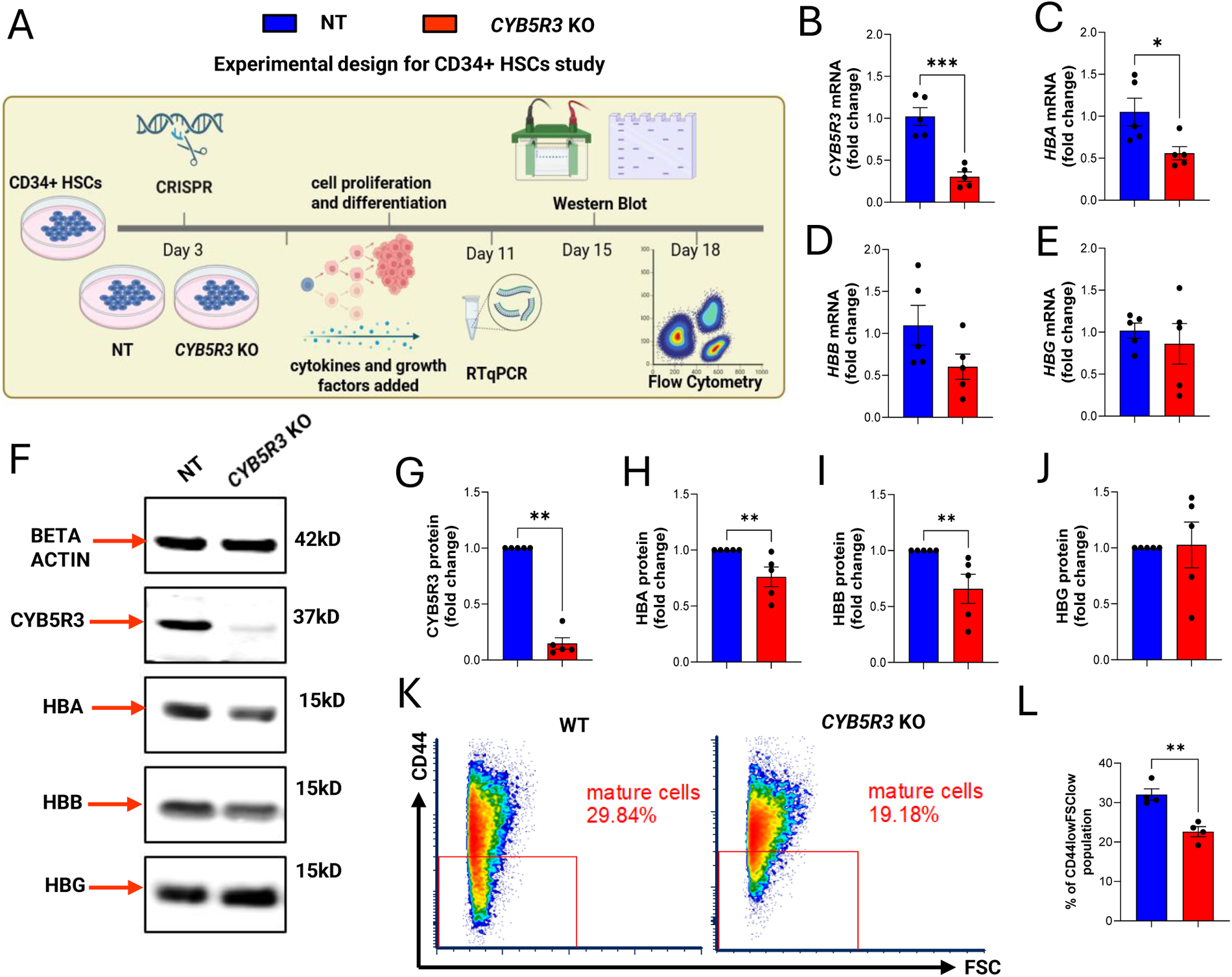
Knocking out CYB5R3 in CD34+ HSCs lowers globin level and hinders erythroblast maturation. **A.** Experimental design for CD34+ HSC study. **B-E.** Quantification of CYB5R3 and globin mRNA levels on day 11. **F.** Representative blots showing the expression of CYB5R3 and globin proteins on day 15. The blots were cut to separate CYB5R3 and Beta Actin, and the globins, due to globins having very high-intensity bands. Each type of globin was stained and quantified, then the blot was stripped to stain for the others separately, since they are of the same size. **G-J.** Quantification of CYB5R3 and globin proteins on day 15. **K.** Representative flow cytometry plots showing CD44_low_FSC_low_ population on day 18. **L.** Quantification of the percentage of CD44l_ow_FSC_low_ population on day 18. Flow cytometry was performed using a BD Biosciences LSR II. Data were analyzed using FCS Express 7 Research Edition (De Novo Software). GraphPad Prism version 10 was used for statistical analyses. Data are stated as mean ± standard error of the mean unless indicated otherwise. Parametric and non-parametric data with 1 variable were analyzed using Student’s t-test and Mann-Whitney test, respectively. F-test was used to compare variances. Grouped data with 2 variables were analyzed using 2-way ANOVA. * indicates p<0.05 and was considered significant. CYB5R3: Cytochrome B5 reductase 3; NT: Non-targeting, KO: Knockout; HBA: Alpha Globin; HBB: Beta Globin; HBG: Gamma globin

### Absence of hematopoietic lineage-specific CYB5R3 lowers the erythropoietic induction by hypoxia in male mice through an erythroblast differentiation defect

Next, we induced erythropoiesis in mice by exposing them to chronic hypoxia for 3 weeks **(Figure 1A)**. Male *CYB5R3* KO mice, when compared to the male WTs, exhibited a significantly lower induction of erythrocytes (25.17% vs. 38.69%), hemoglobin (32.77% vs. 43.29%), and hematocrit (43.34% vs. 54.20%) **(Figure 1B)**. However, no significant differences were observed between the groups for platelets, WBCs, or spleen size **(Supp. Figures 4A-C),** regardless of sex. The Stage II (32% higher), and Stage III and Stage IV (60-80% lower) erythroblast populations were significantly different in the male *CYB5R3* KO versus WTs **(Figures 1C-G)**. There was no notable variation in the HSCs, CMPs, MEPs, or GMPs between the two groups of mice, regardless of sex **(Supp. Figures 4D-I)**. This indicated that the reduced erythropoietic induction in male *CYB5R3* KO mice, under hypoxic stress, was due to a lower percentage of late-stage erythroblasts in the bone marrow.

**Figure 4.**
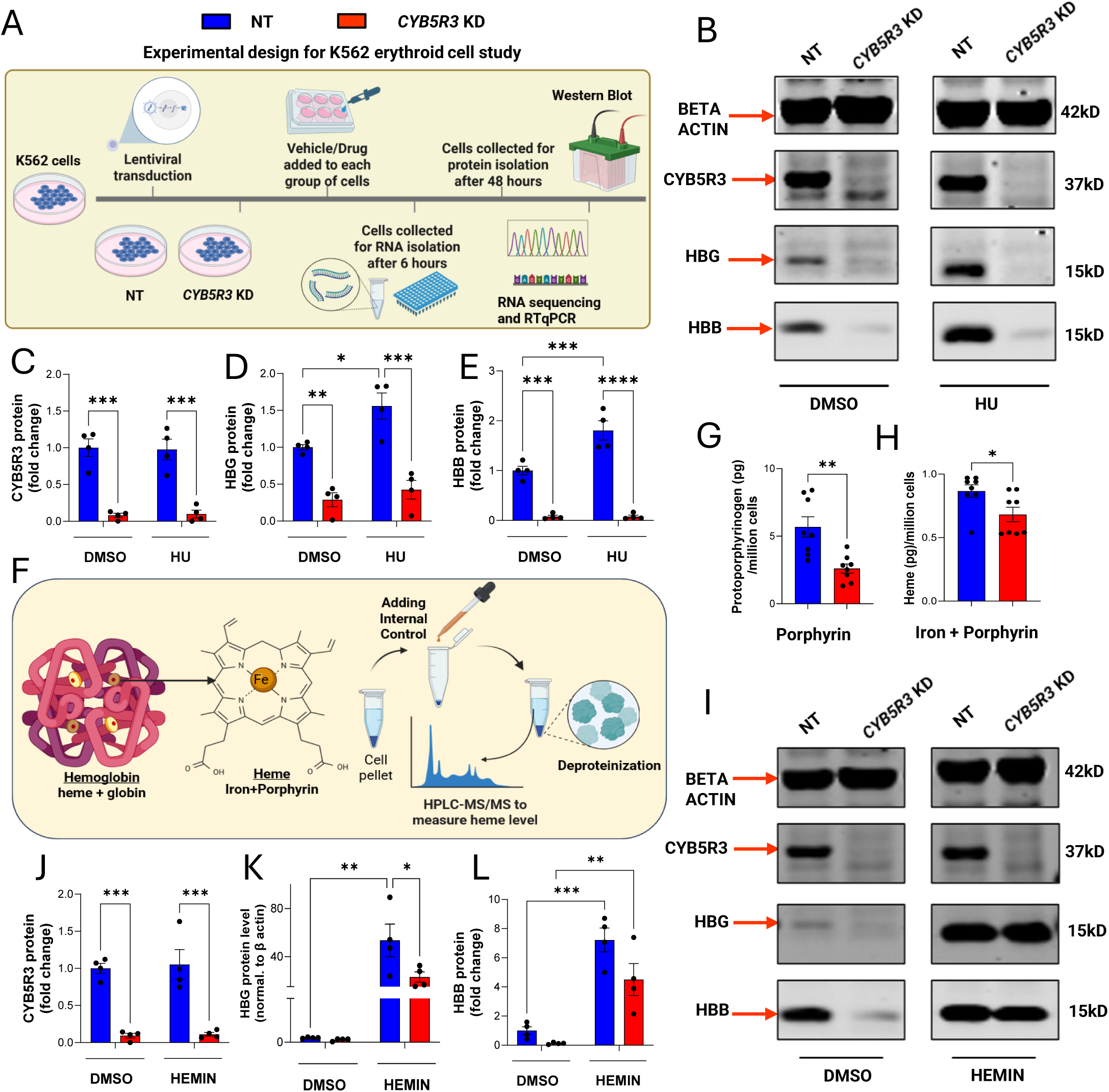
Knocking down CYB5R3 in K562 cells lowers the globin level, which cannot be rescued by hydroxyurea due to heme deficiency. **A.** Experimental design for K562 cells. **B.** Representative blots showing the expression of CYB5R3 and globin proteins after 48 hours of treatment with DMSO or HU. Each type of globin was blotted separately since they are of the same size. **C-E.** Quantification of CYB5R3 and globin proteins after 48 hours of treatment with DMSO or HU. **F.** Schematic diagram showing the components of hemoglobin and heme. **G-H.** HPLC-MS/MS quantification of protoporphyrinogen IX (**G**, MRM 563.3/504.4) and heme (**H**, MRM 616.1/557.1). **I.** Representative blots showing the expression of CYB5R3 and globin proteins after 48 hours of treatment with DMSO or hemin. Each type of globin was blotted separately since they are of the same size. **J-L.** Quantification of CYB5R3 and globin proteins after 48 hours of treatment with DMSO or hemin. GraphPad Prism version 10 was used for statistical analyses. Data are stated as mean ± standard error of the mean unless indicated otherwise. Parametric and non-parametric data with 1 variable were analyzed using Student’s t-test and Mann-Whitney test, respectively. F-test was used to compare variances. Grouped data with 2 variables were analyzed using 2-way ANOVA. * indicates p<0.05 and was considered significant. CYB5R3: Cytochrome B5 reductase 3; NT: Non-targeting, KD: Knockdown; HBB: Beta Globin; HBG: Gamma globin; HU: Hydroxyurea; HPLC-MS/MS: HighPerformance Liquid Chromatography-Mass-Spectrometry

### Splenectomy induces a significant drop in erythropoiesis in CYB5R3 KO mice, which further impacts erythropoietic induction by hypoxia, specifically in male mice

To remove the splenic contribution during stress erythropoiesis induced by hypoxia, we performed splenectomy **(Figure 2A)**. 8 weeks after splenectomy, a significant drop in the erythrocyte parameters was detected in *CYB5R3* KO mice compared to WT mice: erythrocytes (-3.974% vs. 5.553%), hemoglobin (-4.459% vs 3.693%), hematocrit (-4.025% vs. 4.194%) **(Figure 2B)**. Splenectomy induced thrombocytosis and leukocytosis in both groups, as seen previously in human patients **(Supp. Figures 5A-B)** ^27^. We then exposed the splenectomized mice to chronic hypoxia and remeasured the hematological parameters **(Figure 2A)**. An overall lower induction of erythrocytes (14.37% vs. 24.21%), hemoglobin (20.33% vs 32.84%), hematocrit (27.60% vs. 41.98%) **(Figure 2C)**, and a higher induction of platelets (26.25% vs. 3.405%, **Supp. Figures 5C**) were observed in *CYB5R3* KO relative to WT. Unlike the female mice, male-specific analysis revealed significant differences between male *CYB5R3* KO mice and WT mice in the induction of erythrocytes (6.207% vs. 25.40%), hemoglobin (14.93% vs. 38.61%), hematocrit (23.30% vs. 49.17%) **(Figure 2C),** and platelets (29.66% vs. -15.55%, **Supp. Figures 5C**). WBCs did not show any significant differences between the groups, regardless of sex **(Supp. Figure 5D)**. In the bone marrow, the Stage IV erythroblast population was 63% **(Figure 2D-H)**, and the HSC population was 49.67% lower in the male *CYB5R3* KO mice, when compared to male WT mice **(Supp. Figure 5E-F)**. CMPs, MEPs, and GMPs did not differ between the 2 groups, regardless of sex **(Supp. Figure 5G-J)**. Combined, the data showed that hypoxia-induced erythropoietic induction after splenectomy was also diminished in *CYB5R3* KO mice due to erythroblast differentiation defect.

**Figure 5.**
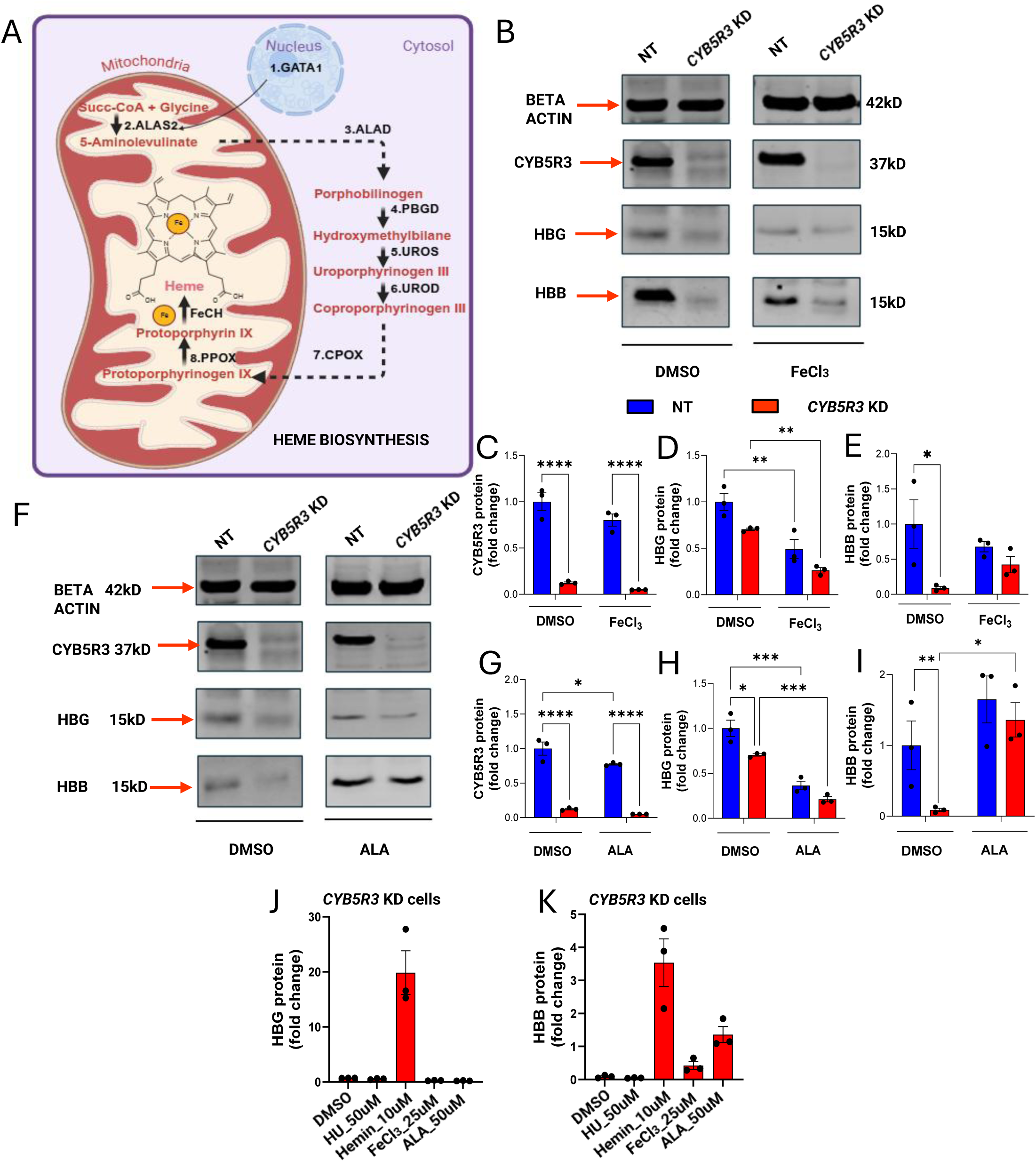
Globin depletion due to heme deficiency cannot be rescued entirely by iron or a porphyrin intermediate. **A.** Heme biosynthesis pathway. **B** Representative blots showing the expression of CYB5R3 and globin proteins after 48 hours of treatment with DMSO or FeCl_3_. Each type of globin was blotted separately since they are of the same size. **C-E.** Quantification of CYB5R3 and globin proteins after 48 hours of treatment with DMSO or FeCl_3_. **F.** Representative blots showing the expression of CYB5R3 and globin proteins after 48 hours of treatment with DMSO or ALA. Each type of globin was blotted separately since they are of the same size. **G-I.** Quantification of CYB5R3 and globin proteins after 48 hours of treatment with vehicle or ALA. **J.** Plot comparing the effect of each treatment on the HBG protein level. **K.** Plot comparing the effect of each treatment on the HBB protein level. GraphPad Prism version 10 was used for statistical analyses. Data are stated as mean ± standard error of the mean unless indicated otherwise. Parametric and non-parametric data with 1 variable were analyzed using Student’s t-test and Mann-Whitney test, respectively. F-test was used to compare variances. Grouped data with 2 variables were analyzed using 2-way ANOVA. * indicates p<0.05 and was considered significant. GATA1: GATA binding factor 1; succinyl Co-A: Succinyl-coenzyme A; ALAD: ALA dehydratase; PBGD: Porphobilinogen deaminase; UROS: Uroporphyrinogen cosynthetase; UROD: Uroporphyrinogen decarboxylase; CPOX: coproporphyrinogen III oxidase; PPOX: Protoporphyrinogen III oxidase; FECH: ferrochelatase; CYB5R3: Cytochrome B5 reductase 3; NT: Non-targeting; KD: Knockdown; HBB: Beta Globin; HBG: Gamma globin; FeCl_3_: Ferric Chloride; ALA: 5-Aminolevulinic acid

### Splenectomy exacerbates the CYB5R3 knockout effect on stress erythropoiesis, more severely in male mice

Splenectomy reduced the hypoxia-induced erythropoiesis in *CYB5R3* KO mice significantly, relative to hypoxia alone **(Supp. Figures 6A-C)**; erythrocytes (overall 14.37% vs. 28.14%, male only 6.207% vs. 25.17%), hemoglobin (overall 20.33% vs. 33.08%, male only 14.93% vs. 32.77%), and hematocrit (overall 27.60% vs. 43.04%, male only 23.30% vs. 43.34%). Hypoxia after splenectomy also increased thrombocytosis and leukocytosis in both groups substantially, due to the lack of splenic sequestration **(Supp. Figures 6D-E)**; however, platelet induction was higher in *CYB5R3* KO mice than WT mice. Hence, it can be concluded that the removal of the spleen significantly affected hypoxia-induced erythropoiesis in *CYB5R3* KO mice, with a more severe effect in males.

**Figure 6:**
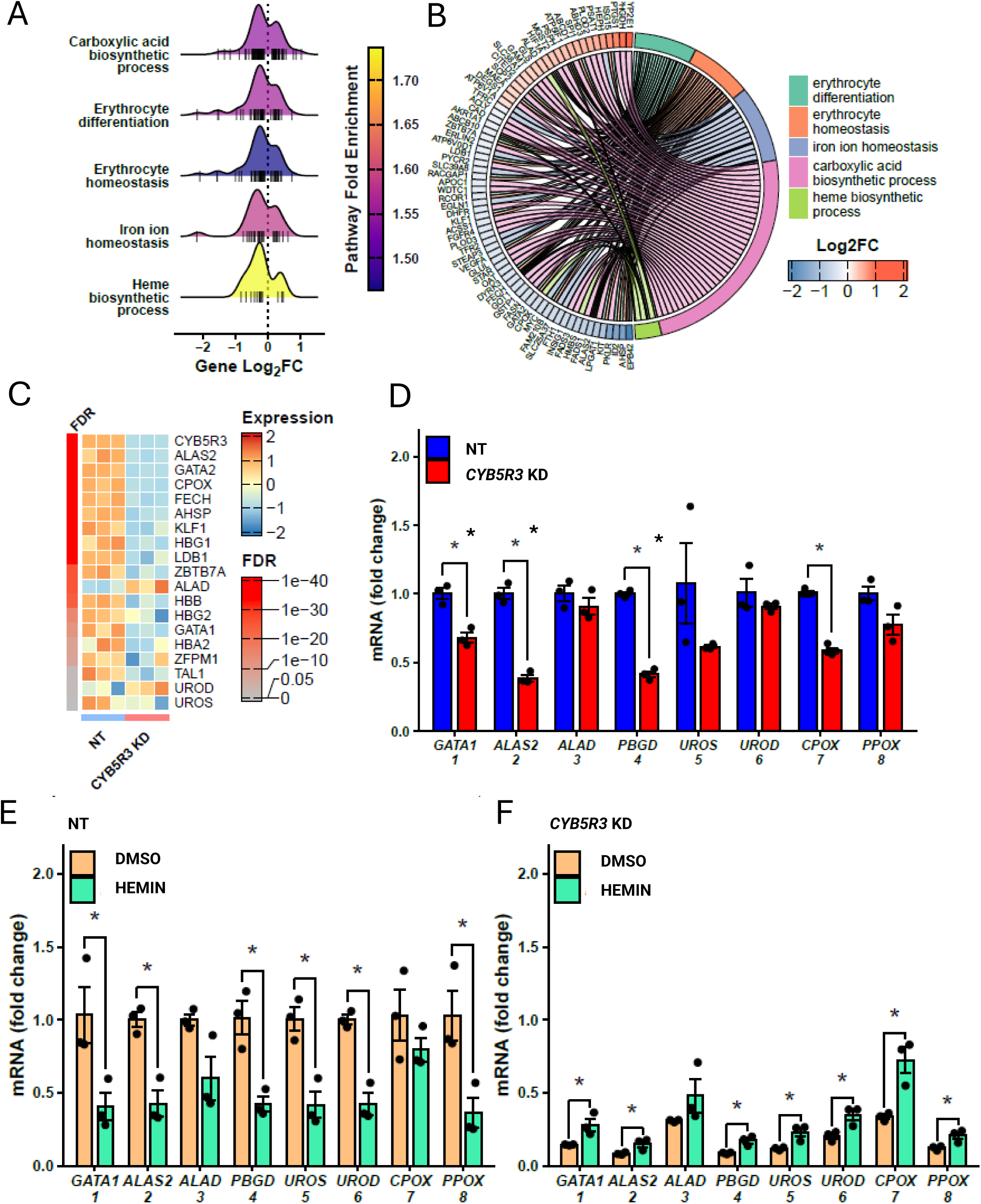
Regulatory factors maintaining heme biosynthesis and erythropoiesis are suppressed in CYB5R3 KD K562 cells. Bulk RNA sequencing analysis shows the transcriptomic differences between NT and *CYB5R3* KD K562 cells. Differentially expressed genes, comparing CYB5R3 KD to NT, with a false discovery rate (FDR)<0.05, are used in an overrepresentation test to identify significantly enriched pathways. **A.** Significantly enriched Gene Ontology biological processes (p<0.05) related to hematopoiesis are included in the density plot. The height of the density plot indicates the number of genes in a pathway, with the log2-transformed fold changes (log2FC, CYB5R3 KD vs NT) indicated on the x-axis. Individual genes involved in the pathway are shown as bars under the density plot. The color of the density plot represents the pathway fold enrichment in the over-representation test. **B.** The chord plot maps the top differentially expressed genes (FDR<0.001) to the aforementioned Gene Ontology pathways. **C.** The heatmap shows the relative expression (z-scores) of erythroid- and heme-biosynthesis-related genes. The left color bar indicates the gene’s FDR in the differential expression analysis. **D.** Quantification of the mRNA levels of *GATA1* and the enzymes (1-8) involved in the heme biosynthesis pathway in independent experiments using NT and *CYB5R3* KD K562 cells. **E.** Quantification of the mRNA levels of GATA1 and the enzymes (1-8) involved in the heme biosynthesis pathway in NT cells after hemin treatment. **F.** Quantification of the mRNA levels of *GATA1* and the enzymes (1-8) involved in the heme biosynthesis pathway in *CYB5R3* KD cells after hemin treatment. * indicates p<0.05 using Student’s t-test and was considered significant. CYB5R3: Cytochrome B5 reductase 3; NT: Non-targeting; KD: Knockdown; GATA1: GATA binding factor 1; ALAS2: 5-aminolevulinic acid synthase 2; ALAD: ALA dehydratase; PBGD: Porphobilinogen deaminase; UROS: Uroporphyrinogen III synthase; UROD: Uroporphyrinogen III decarboxylase; CPOX: Coproporphyrinogen III oxidase; PPOX: Protoporphyrinogen III oxidase

### Knocking out CYB5R3 in CD34+ HSCs lowers globin level and hinders erythroblast maturation

To assess the role of CYB5R3, we induced erythropoiesis in CD34+ HSCs and collected samples at different time points for analysis **(Figure 3A).** Expression of *CYB5R3* mRNA on day 11 (before organelle ejection is initiated, **Figure 3B**) and CYB5R3 protein on day 15 **(Figures 3F-G)** in NT cells vs. *CYB5R3* KO cells confirmed its deletion in the KO group. Since hemoglobin formation is the key step of erythropoiesis, we measured mRNA **(Figures 3C-E)** and protein expression **(Figures 3F, 3H-J)** for alpha globin (HBA), beta globin (HBB), and gamma globin (HBG). mRNA level for HBA and protein level for HBA and HBB were significantly lower in *CYB5R3* KO cells. A smaller percentage of enucleated reticulocytes and mature erythrocytes (CD44_low_FSC_low_ cells)^19^ in *CYB5R3* KO cells (**Figures 3K-L**), determined by flow cytometric analysis on day 18, demonstrated diminished erythrocyte differentiation. Together, the data suggested that globin level and erythropoiesis were affected in CD34+ HSCs due to CYB5R3 deficiency.

### Knocking down CYB5R3 in K562 cells lowers the globin level, which cannot be rescued by HU due to heme deficiency

We continued our investigation using K562 cells for an in-depth understanding of the mechanistic role of CYB5R3 in erythropoiesis **(Figure 4A)**. The mRNA level **(Supp. Figure 7A)** and protein expression **(Figures 4B-C)** confirmed CYB5R3 deletion in *CYB5R3* KD K562 cell line. As published previously, the K562 cells did not express HBA.^28,29^ Although the baseline mRNA changes (with vehicle, DMSO) were not significant **(Supp. Figure 7B-C)**, CYB5R3 deficiency depleted the baseline protein levels of HBB and HBG to less than 50% **(Figures 4B, D-E)** of NT cells. HU, an erythroid differentiation-inducing agent, triggered a 1.5-fold globin induction in NT cells relative to baseline. By contrast, HU did not induce globin accumulation in *CYB5R3* KD cells at either the mRNA **(Supp. Figures 7B-C)** or the protein level **(Figures 4B, D-E)**. The data were consistent with those obtained using *CYB5R3* KO CD34+ HSCs.

**Figure 7:**
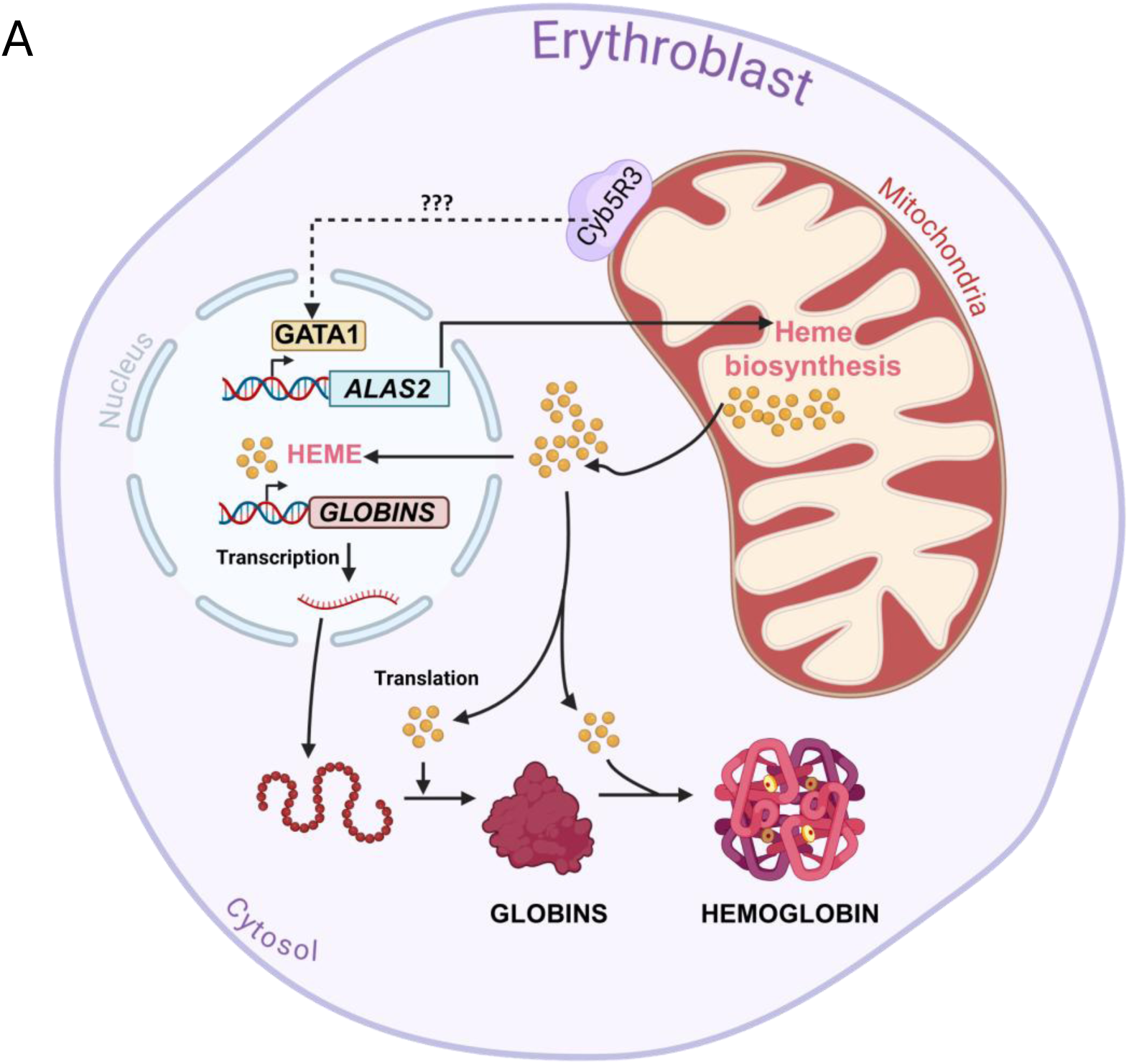
Proposed mechanistic role of CYB5R3 in erythropoiesis. Based on our data, CYB5R3 regulates the expression of erythroid-specific genes involved in heme biosynthesis, including GATA binding factor 1 (GATA1) and 5-aminolevulinic acid synthase 2 (ALAS2). ALAS2 is the rate-limiting enzyme in the heme biosynthesis pathway. Without sufficient ALAS2, the rest of the heme biosynthesis process is not induced; the enzymes downstream are suppressed, resulting in heme deficiency. The absence of heme negatively affects the transcription and translation of globins. Additionally, insufficient heme may affect the stability of globins, as heme and globin together form a stable hemoglobin molecule with native folding. However, whether CYB5R3 carries out these actions directly or through mediators remains to be determined.

Since globin expression is regulated by heme, we measured levels of the heme intermediate, protoporphyrinogen IX, and the final product, heme, using high-performance liquid chromatography-mass spectrometry (HPLC-MS/MS) **(Figure 4F)** ^30,31,32^. We found that both the protoporphyrinogen and heme levels were reduced by approximately 40% in *CYB5R3* KD cells relative to NT **(Figures 4G-H)**. Hemin, a commercially available form of oxidized heme, was then supplied to rescue globin formation in *CYB5R3* KD cells. While no effect was observed on the mRNA levels of CYB5R3, HBB, or HBG **(Supp. Figures 7D-F)**, the HBB and HBG proteins were substantially rescued, increasing approximately 30-40 times relative to the vehicle-treated *CYB5R3* KD cells **(Figures 4I-L)**. Together, these findings confirm that the observed globin depletion is happening due to heme deficiency in *CYB5R3* KD K562 cells.

### Globin depletion due to heme deficiency cannot be rescued entirely by iron or a porphyrin intermediate

Heme biosynthesis is a multi-step process involving multiple enzymes **(Figure 5A)**. Since iron is an essential component of heme and CYB5R3 is associated with heme reduction, we supplied iron as FeCl_3_ to test if *CYB5R3* KD cells **(Supp. Figure 8A**, **Figures 5B-C)** were iron-deficient^33,34,11^. We found that FeCl_3_ did not rescue the globin mRNA levels **(Supp. Figures 8B-C)**; however, the HBB protein expression trended upward **(Figures 5B, D-E)**. Subsequently, in consideration of a possible role for deficient synthesis of heme substrates, we supplied *CYB5R3* KD cells **(Supp. Figure 8D**, **Figures 5F-G)** with 5-aminolevulinic acid (ALA), a porphyrin precursor synthesized in the first and rate-limiting step of the heme biosynthesis pathway^8^. Although mRNA levels were not positively affected **(Supp. Figures 8E-F)**, HBB protein expression also trended upward with ALA supplementation **(Figures 5F, H-I)**. When we combined the changes observed after each treatment in KD cells into a graph **(Figures 5J-K)**, it became apparent that hemin at the lowest of the treatment doses caused the highest increase in globin levels. Although FeCl_3_ and ALA partially rescued the HBB protein, the opposite occurred with the HBG protein. These findings suggested that the deficiency of iron or porphyrin precursors was not the actual underlying cause of heme deficiency.

### Regulatory factors maintaining heme biosynthesis and erythropoiesis are suppressed in CYB5R3 KD K562 cells

Because FeCl_3_ and ALA supplementation both failed to fully restore globin expression, we reasoned that the downstream heme biosynthesis pathway may be compromised. Bulk RNA sequencing revealed that differentially expressed genes (DEGs) (false discovery rate < 0.05), due to the loss of CYB5R3, are enriched in erythropoiesis-related Gene Ontology (GO) pathways, including carboxylic acid biosynthesis, erythrocyte differentiation, erythrocyte homeostasis, iron ion homeostasis, and the heme biosynthetic process (**Figure 6A**). Within these pathways, the majority of DEGs were downregulated in *CYB5R3* KD K562 cells (**Figure 6A**). Notably, among the most significant DEGs (FDR < 0.001), aminolevulinic acid synthase 2 (*ALAS2*), coproporphyrinogen III oxidase (*CPOX*), LIM-domain-binding protein 1 (*LDB1*), krueppel-like factor 1 (*KLF1*), ferrochelatase (*FeCH*), and GATA-binding factor 2 (*GATA2*) are downregulated in CYB5R3 KD cells (**Figure 6B**). Additionally, the expression of *HBG1*, *HBG2*, *HBB*, and GATA-binding factor 1 (*GATA1*) were also suppressed by CYB5R3 silencing (**Figure 6C**). The aforementioned genes are critical for the regulation and maintenance of heme biosynthesis and erythropoiesis, and their downregulation indicated a defect in heme biosynthesis.

We then quantified the mRNA levels of *GATA1,* a master transcription factor^35–37^, and enzymes that are involved in the heme biosynthetic pathway: *ALAS2,* aminolevulinic acid dehydratase *(ALAD),* porphobilinogen deaminase *(PBGD),* uroporphyrinogen synthase III synthase (*UROS),* uroporphyrinogen decarboxylase *(UROD), CPOX,* and protoporphyrinogen IX oxidase (*PPOX)* ^32,38^. All of these genes were significantly down-regulated in the *CYB5R3* KD group relative to NT (**Figure 6D**). When supplemented with hemin, mRNA levels of these genes decreased in NT cells (**Figure 6E**), while a modest but significant increase was noted in *CYB5R3* KD K562 cells (**Figure 6F**). Taken together, these data suggested that knocking down *CYB5R3* downregulated the genes associated with heme biosynthesis and erythropoiesis, leading to heme deficiency-induced globin depletion that can be rescued by hemin supplementation.

## Discussion

According to the World Health Organization (WHO), anemia caused a loss of 50 million years of healthy life in 2019 due to disability^1^. The primary reasons are malnutrition, chronic diseases, menstruation, inflammation, or pregnancy, all of which make a person susceptible to oxidative stress^1,3,39^. Prior work showed that oxidative stress reduces the level of erythropoietin secreted by the kidney for inducing erythropoiesis^4^. Although the lifespan of erythrocytes is around 120 days, if faced with oxidative injury in the circulation, suicidal death ensues^3^. Erythrocytes are particularly susceptible to oxidative stress due to their high content of heme and polyunsaturated fatty acids, as well as their primary role in transporting oxygen^39^. As such, they possess a stringent redox regulatory mechanism, in which CYB5R3 has a crucial contribution^11,39^. Our previous study, which utilized both clinical and *in vitro* data, strongly suggested a role for CYB5R3 in HU-induced HbF formation and erythrocyte maturation^19^. However, in this study, we have demonstrated for the first time that CYB5R3 plays an additional role in controlling erythrocyte development by regulating heme biosynthesis.

We have used hematopoietic lineage-specific *CYB5R3* KO mice to determine the role of CYB5R3 in erythropoiesis. At steady state, although MEPs were higher in the KO mice, this did not result in significant changes in blood parameters. The KO mice had a darker blood color due to the deficiency of CYB5R3 as a met-hemoglobin reductase. We next induced erythropoiesis using hypoxia. Hypoxia is a state of low oxygenation that triggers the kidney to produce more erythropoietin, leading to increased erythropoiesis for restoring homeostasis^40,41^. The spleen, which is the primary extramedullary site of stress erythropoiesis, and bone marrow both participate in responding to hypoxic stress^42,43^. In *CYB5R3* KO mice, hypoxia-stimulated erythropoiesis was significantly altered, which was further amplified in the absence of the spleen and more pronounced in males. Erythroblasts are differentiated from MEPs, which in turn are derived from HSCs^44,45^. Since the MEP populations were similar between groups after hypoxia or splenectomy-hypoxia combination, the observed lower percentage of late-stage erythroblasts in the bone marrow was attributed to reduced erythropoietic induction. This reasoning suggests a defect in the terminal differentiation pathway, wherein erythroblasts are unable to progress beyond the early stages to become mature erythrocytes. Additionally, splenectomy resulted in more rapid exhaustion of HSCs in male *CYB5R3* KO mice under hypoxic conditions in the bone marrow. As for the overall greater impact of *CYB5R3* KO on male mice, we recently showed that cardiomyocyte-specific *CYB5R3* KO causes cardiac hypertrophy, bradycardia, and sudden cardiac death in males, but not in females^13^. A previous study suggests that in humans, females recover respiratory functions faster than males after hypoxic exposure^46^. It was also shown that female rats have a higher hypoxic and hypercapnic ventilatory response than male rats^47^. In SCD, where stress erythropoiesis is a common occurrence, males have increased morbidity and lower survival relative to females^48,49^. These sex differences are generally attributed to estrogen levels, which provide females with a greater overall protection against hypoxia^47,50^.

To investigate the cause of the observed erythroblast differentiation defect in the animal model, we initiated an *in vitro* study using erythroid cells. After inducing erythropoiesis in CD34+ HSCs and K562 cells **(Figures 3-4)**, we found significantly lower total hemoglobin in *CYB5R3* KO cells compared to NT control. Even HU, a commonly used erythroid differentiation-inducing agent^51^ and the treatment of choice for patients with SCD, could not cause adult hemoglobin (HBB) or fetal hemoglobin (HbF/HBG) accumulation in CYB5R3-deficient K562 cells^52,53^. This was in agreement with our previous study showing the inability of HU to significantly induce F cells in *CYB5R3* KO CD34+ HSCs ^19^. HPLC-MS/MS showed heme deficiency occurred alongside globin depletion in *CYB5R3* KD K562 cells. Heme/hemin, a hemoglobin-inducing agent, as seen in previous studies, substantially rescued globin expression in *CYB5R3* KD K562 cells^54–56^. It is well-established that heme inhibits the repressor BTB-domain and CNC homology 1 (BACH1) and activates the inducer nuclear factor erythroid 2-related factor 2 (NRF2), thereby activating globin transcription^30,57^. Heme also inhibits heme-regulated-eIF2α kinase (HRI) and activates the translation initiation factor alpha subunit of eukaryotic initiation factor 2 (eIF2α) for globin protein synthesis^58^. Under heme deficiency, there is not enough heme available to inhibit HRI or to be incorporated within the globins to form stable hemoglobin molecules^6,59^. Hence, HU-induced hemoglobin transcription is by itself insufficient to support hemoglobin formation^17^.

Heme contains an iron embedded in a porphyrin ring^34,60^. Given that our cells showed hemin-induced hemoglobin accumulation, we supplemented *CYB5R3* KD cells with iron or porphyrin precursor ALA **(Figure 5)**. Surprisingly, neither iron nor ALA could rescue hemoglobin expression like hemin. A genomic screening using NT and *CYB5R3* KD K562 cells **(Figure 6)** revealed differential expression of genes connected to pathways associated with erythropoiesis and heme biosynthesis. In CYB5R3 deficiency, we identified downregulation of essential erythropoiesis-associated genes: *LDB1, KLF1, GATA2, GATA1*; and heme biosynthesis-associated genes: *ALAS2, CPOX,* and *FeCH*. GATA2 is known to directly activate GATA1 expression in early erythroid progenitors^37,61^, followed by GATA1 accelerating its own expression in the later stages^37^. GATA1 binds co-activators such as LDB1 and T-cell acute lymphocytic leukemia protein 1 (TAL1) to activate erythropoiesis^62–64^. Additionally, it recruits co-repressors such as zinc finger protein (ZFPM1) to repress target genes that inhibit erythropoiesis^62,64^. GATA1 also induces the expression of ALAS2, which is required for heme biosynthesis at the later stages of erythroblast differentiation^8^. Deficiency of KLF1 has been reported to cause impaired enucleation in orthochromatic erythroblasts^65^. Moreover, decreased levels of mRNA for *GATA1* and heme biosynthesis enzymes, quantified using real-time qPCR, in *CYB5R3* KD K562 cells were consistent with RNA-seq findings. Although additional heme modestly induced enzyme transcript levels in *CYB5R3* KD cells, it had the opposite effect in NT cells. This suggests that excess heme in wild-type cells exerts negative feedback regulation on the heme biosynthesis pathway, whereas an induction is only possible in a state of deficiency^35,36,66^.

Based on our experimental outcomes, we propose a transcriptional role for CYB5R3 in erythropoiesis-associated heme biosynthesis **(Figure 7)**. The transcriptional role indicates that CYB5R3 deficiency downregulates erythroid-associated genes, suppressing the critical transcription factors and enzymes that regulate heme biosynthesis. This, in turn, hinders hemoglobin formation as observed in our *in vitro* study, followed by improper erythroid differentiation and development, as observed in the animal model. These findings support previously published observations, where defects in the heme biosynthesis pathway created developmental issues in erythroid cells, causing erythropoietic disorders, including anemia and porphyria^8,32,66^. These results also help explain the diminished HU-induced HbF response observed in SCD patients carrying the hypomorphic T117S variant of CYB5R3 and diminished HU-induced F cells in *CYB5R3* KO CD34+ HSCs^19^. The most plausible explanation for CYB5R3’s role in erythropoiesis can be its ability to regulate the NO-sGC-cGMP signaling pathway, which warrants further investigation^11,16,17,67^.

A few caveats are noted. We initially used CD34+ HSCs to set the premise and then continued an in-depth analysis using K562 cells, due to the inconvenience we and others faced in using CD34+ cells^25^. The human K562 cell line, like CD34+ HSCs, is a multi-potent hematopoietic precursor cell line that can be differentiated into erythroid cells^54,68^. Although the K562 cell line has been widely used as an erythropoietic model due to its glycophorin A and hemoglobin expression, it has a neoplastic origin and is prone to mutational changes^25^. Additionally, K562 cells do not always express enough HBG, and do not express HBB, ideally^25, 69^. In this case, the K562 cell line expressed enough HBG and HBB at baseline, making it easy to study the mechanistic role of CYB5R3 in erythropoiesis.

In summary, our findings build a foundation for future endeavors to identify CYB5R3-based therapeutics for anemia-targeted intervention. In this study, we have uncovered a crucial second role for CYB5R3 as a regulator of erythropoiesis, the deletion of which causes a male-specific erythroblast differentiation defect because of heme deficiency. These results uncover an unexpected function for CYB5R3 beyond methemoglobin reduction, establishing it as a key metabolic driver of sex-specific stress erythropoiesis and exposing a heme-limited vulnerability that may worsen disease severity in anemia, hemoglobinopathies, and carriers of CYB5R3 loss-of-function variants. These new insights not only refine our understanding of the importance of redox signaling in erythropoiesis but also have the potential to be applied to newer individualized interventions, drug development strategies, and regenerative medicine that specifically target anemia in hemoglobinopathies and other erythropoietic disorders.

## Authorship contributions

FAC performed the experiments, analyzed the data, and wrote the manuscript. ACS designed the research questions, wrote, and edited the manuscript. MS, SAS, MPM, SAH, MK, and SNT assisted with the experiments. SY performed RNA-seq analysis and assisted with the experiments and edits. SRS performed and analyzed the HPLC-MS/MS data and assisted with the edits. KCW and FJS helped with the research questions and edits.

## Data Sharing

Data will be shared upon request to the corresponding author.

