## Supplemental file for "CYB5R3 Controls Sex-Specific Stress Erythropoiesis via Heme-Biosynthesis"

### Supplemental Figure 1

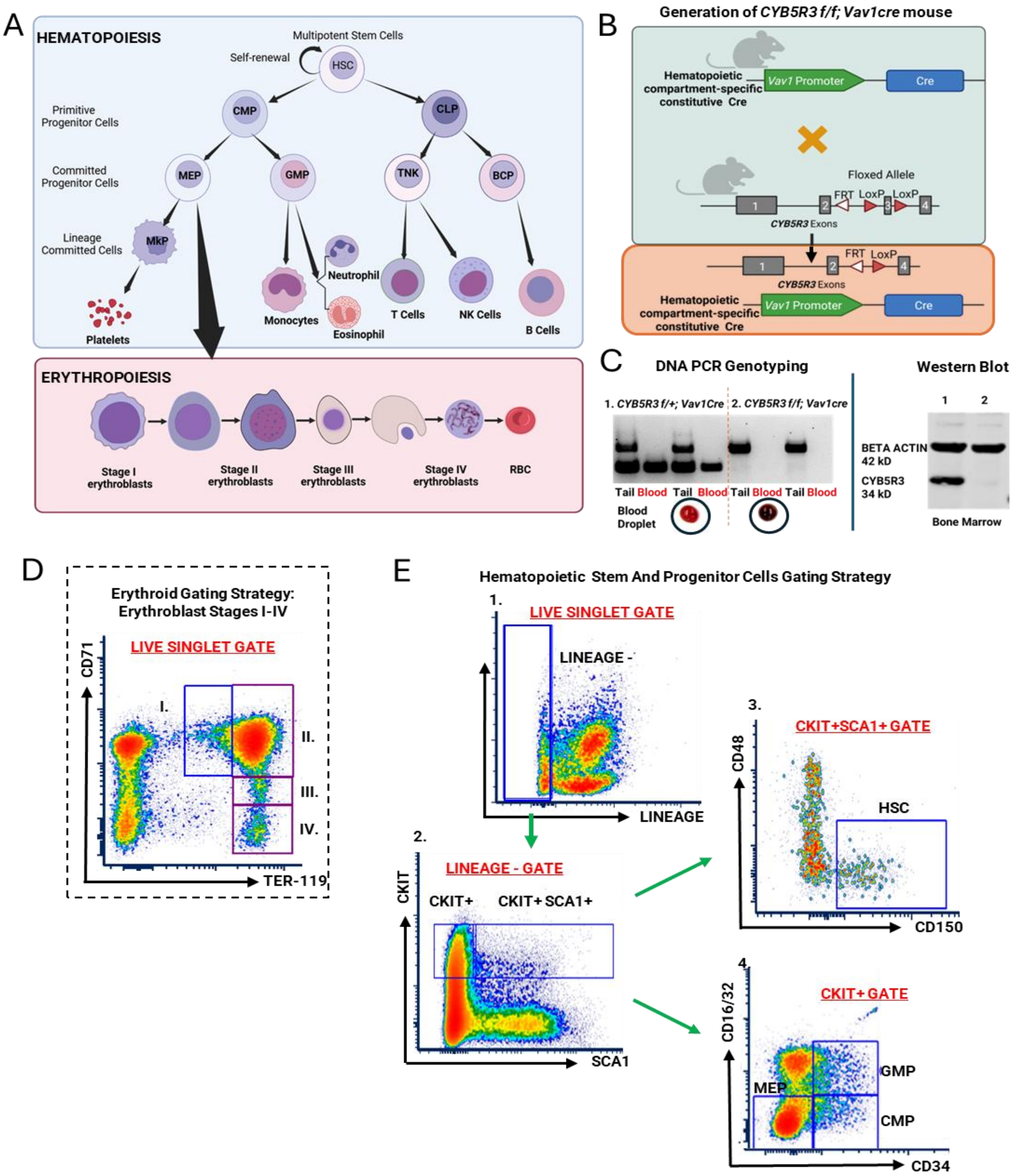


**Generation of mouse and flow cytometry gating strategy**

**A.** Schematic diagram showing hierarchy of hematopoiesis and stages of erythropoiesis. **B.** Generation of hematopoietic lineage-specific *CYB5R3* knockout mouse. **C.** Confirmation of hematopoietic lineage-specific *CYB5R3* knockout in mouse. **D.** Gating strategy for determining different stages of erythroblasts.[1] **E.** Gating strategy for determining HSCs and erythroid progenitors.[2,3] HSC: hematopoietic stem cells; CMP: Common Myeloid Progenitors; CLP: Common Lymphoid Progenitors; MEP: Megakaryocyte Erythroid Progenitors; GMP: Granulocyte Monocyte Progenitors; MkP: Megakaryocyte Progenitors

### Supplemental Figure 2

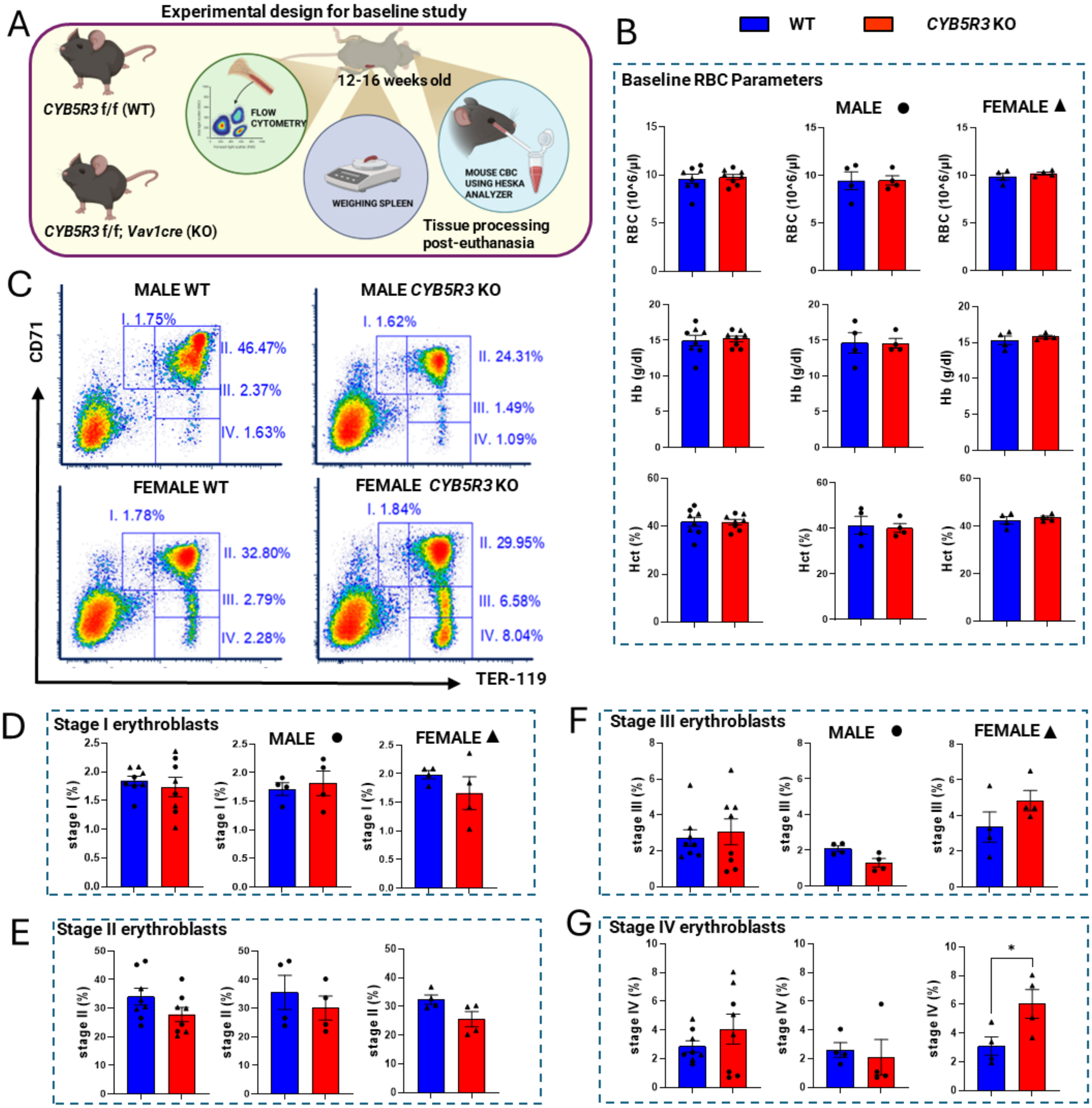


**CYB5R3 in the hematopoietic compartment is dispensable for steady-state erythropoiesis.**

**A.** Experimental design for baseline animal study. **B.** Baseline erythrocyte (RBC) parameters, combined, and males and females separately. **C.** Flow cytometry plots for baseline erythroblast stages in the bone marrow. **D-G.** Quantification of different stages of erythroblasts in the bone marrow at baseline, combined, and males and females separately. Flow cytometry was performed using a BD Biosciences LSR II. Data were analyzed using FCS Express 7 Research Edition (De Novo Software). Data are stated as mean ± standard error of the mean unless indicated otherwise. GraphPad Prism version 10 was used for statistical analyses. Parametric and non-parametric data with 1 variable were analyzed using Student's t-test and Mann-Whitney test, respectively. F-test was used to compare variances. Grouped data with 2 variables were analyzed using 2-way ANOVA. * indicates $p<0.05$ and was considered significant. CYB5R3: Cytochrome B5 reductase 3; WT: Wild-type; KO: Knockout; RBC: red blood cells, erythrocytes; Hb: hemoglobin; Hct: hematocrit

**Supplemental Figure 3**

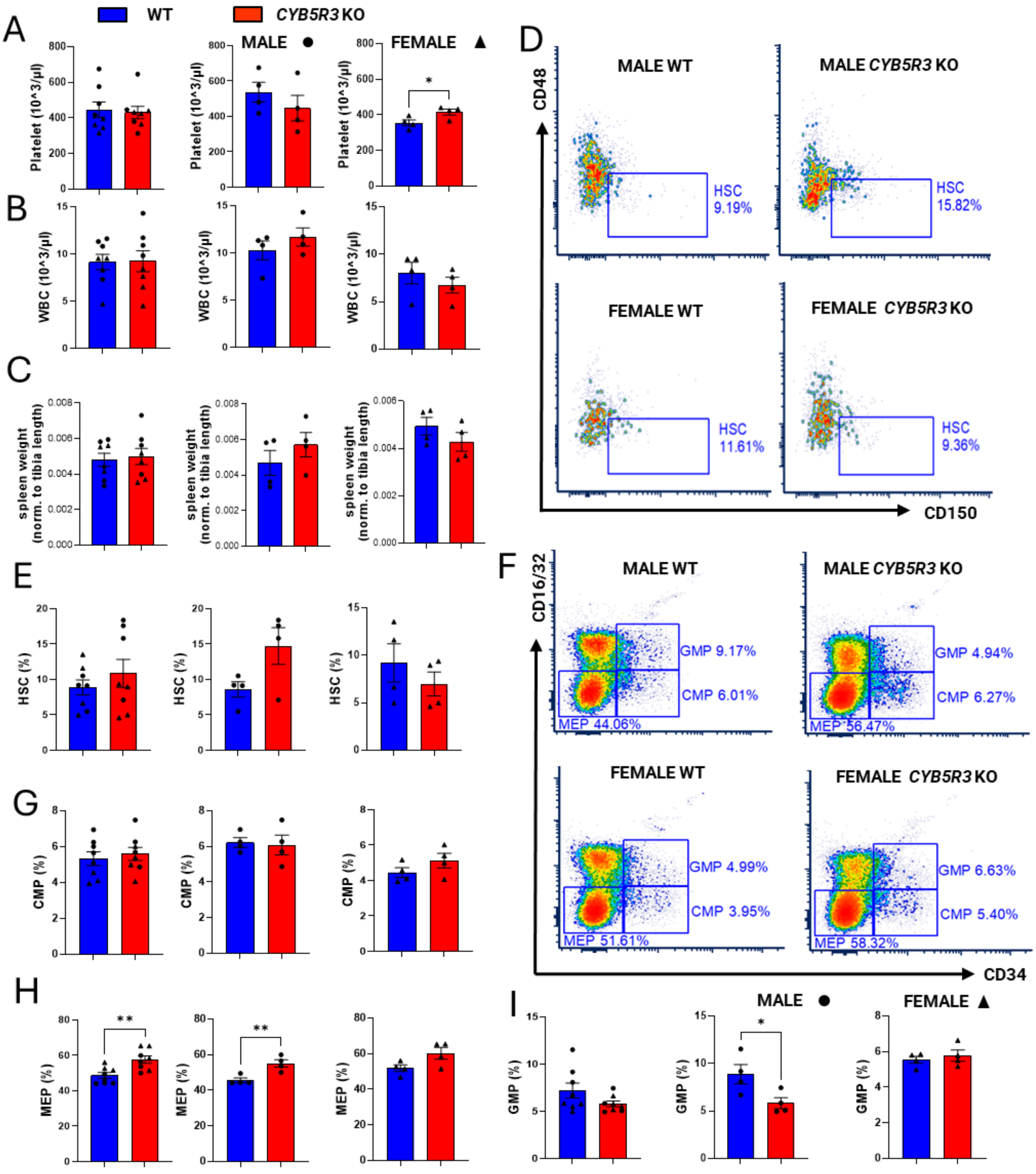


**CYB5R3 in the hematopoietic compartment is dispensable for steady-state erythropoiesis. (extended from Supp. Figure 2)**

**A.** Baseline platelet count, combined, and males and females separately. **B.** Baseline WBC count, combined, and males and females separately. **C.** Normalized spleen weight at baseline, combined, and males and females separately. **D.** Flow cytometry plots showing the percentage of HSC population in the bone marrow at baseline. **E.** Quantification of HSCs in the bone marrow, combined, and males and females separately. **F.** Flow cytometry plots showing the percentage of different types of hematopoietic progenitors in the bone marrow at baseline. **G-I.** Quantification of different types of hematopoietic progenitors in the bone marrow, combined, and males and females separately. Flow cytometry was performed using a BD Biosciences LSR II. Data were analyzed using FCS Express 7 Research Edition (De Novo Software). GraphPad Prism version 10 was used for statistical analyses. Data are stated as mean ± standard error of the mean unless indicated otherwise. Parametric and non-parametric data with 1 variable were analyzed using Student's t-test and Mann-Whitney test, respectively. F-test was used to compare variances. Grouped data with 2 variables were analyzed using 2-way ANOVA. * indicates $p<0.05$ and was considered significant. CYB5R3: Cytochrome B5 reductase 3; WT: Wild-type; KO: Knockout; WBC: white blood cells; HSC: hematopoietic stem cells; CMP: Common Myeloid Progenitors; MEP: Megakaryocyte Erythroid Progenitors; GMP: Granulocyte Monocyte Progenitors

**Supplemental Figure 4**

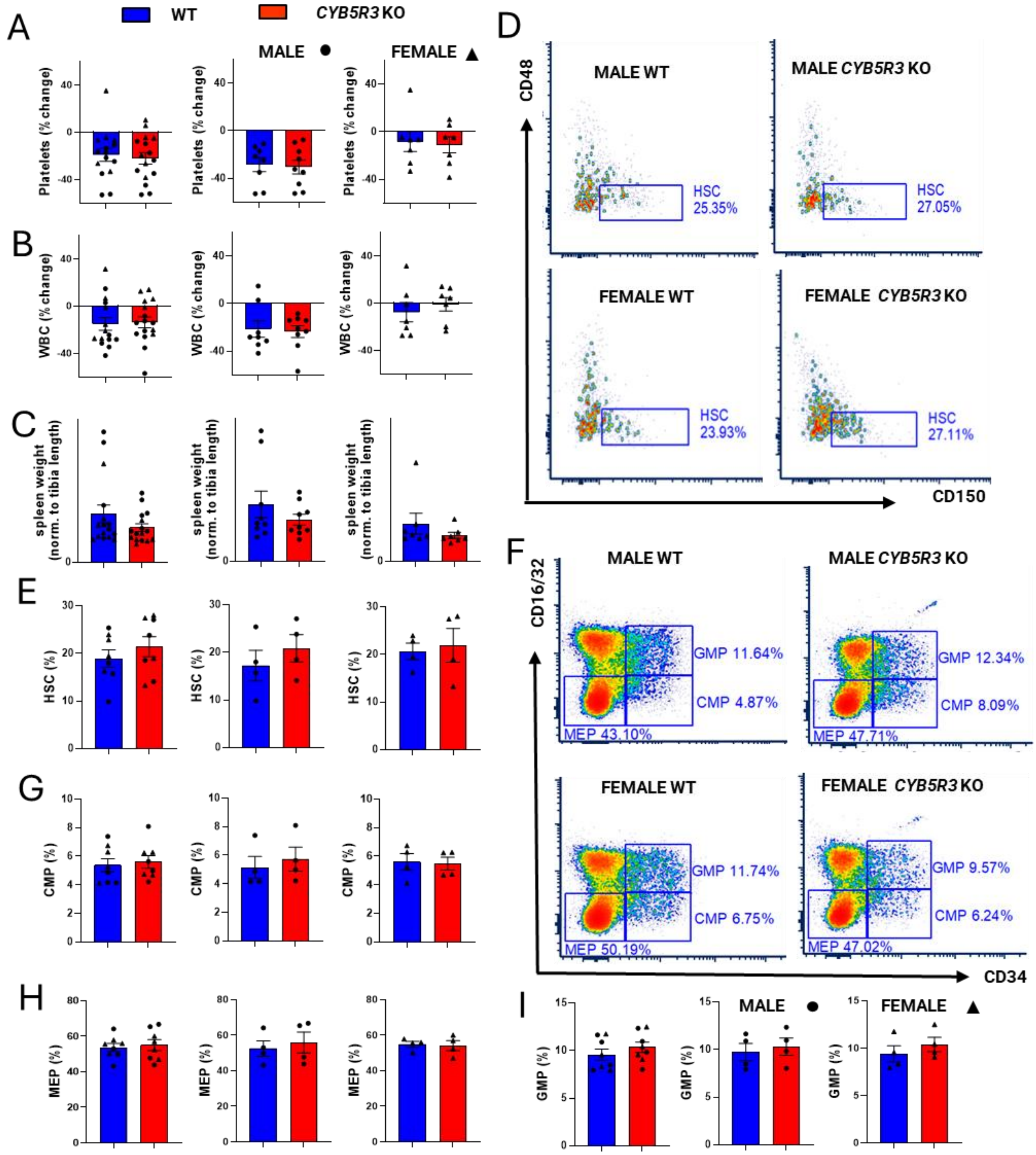


**Absence of hematopoietic lineage-specific CYB5R3 lowers the erythropoietic induction by hypoxia in male mice through an erythroblast differentiation defect.**

**A.** Percentage change in platelets after 3 weeks of hypoxia, combined, and males and females separately. **B.** Percentage change in WBCs after 3 weeks of hypoxia, combined, and males and females separately. **C.** Normalized spleen weight after 3 weeks of hypoxia, combined, and males and females separately. **D.** Flow cytometry plots showing the percentage of HSC population at baseline in the bone marrow after 3 weeks of hypoxia. **E.** Quantification of HSCs in the bone marrow, combined, and males and females separately. **F.** Flow cytometry plots showing the percentage of different types of hematopoietic progenitors in the bone marrow after 3 weeks of hypoxia. **G-I.** Quantification of different types of hematopoietic progenitors in the bone marrow, combined, and males and females separately. Flow cytometry was performed using a BD Biosciences LSR II. Data were analyzed using FCS Express 7 Research Edition (De Novo Software). GraphPad Prism version 10 was used for statistical analyses. Data are stated as mean ± standard error of the mean unless indicated otherwise. Parametric and non-parametric data with 1 variable were analyzed using Student's t-test and Mann-Whitney test, respectively. F-test was used to compare variances. Grouped data with 2 variables were analyzed using 2-way ANOVA. * indicates $p<0.05$ and was considered significant. CYB5R3: Cytochrome B5 reductase 3; WT: Wild-type; KO: Knockout; WBC: white blood cells; HSC: hematopoietic stem cells; CMP: Common Myeloid Progenitors; MEP: Megakaryocyte Erythroid Progenitors; GMP: Granulocyte Monocyte Progenitors

### Supplemental Figure 5

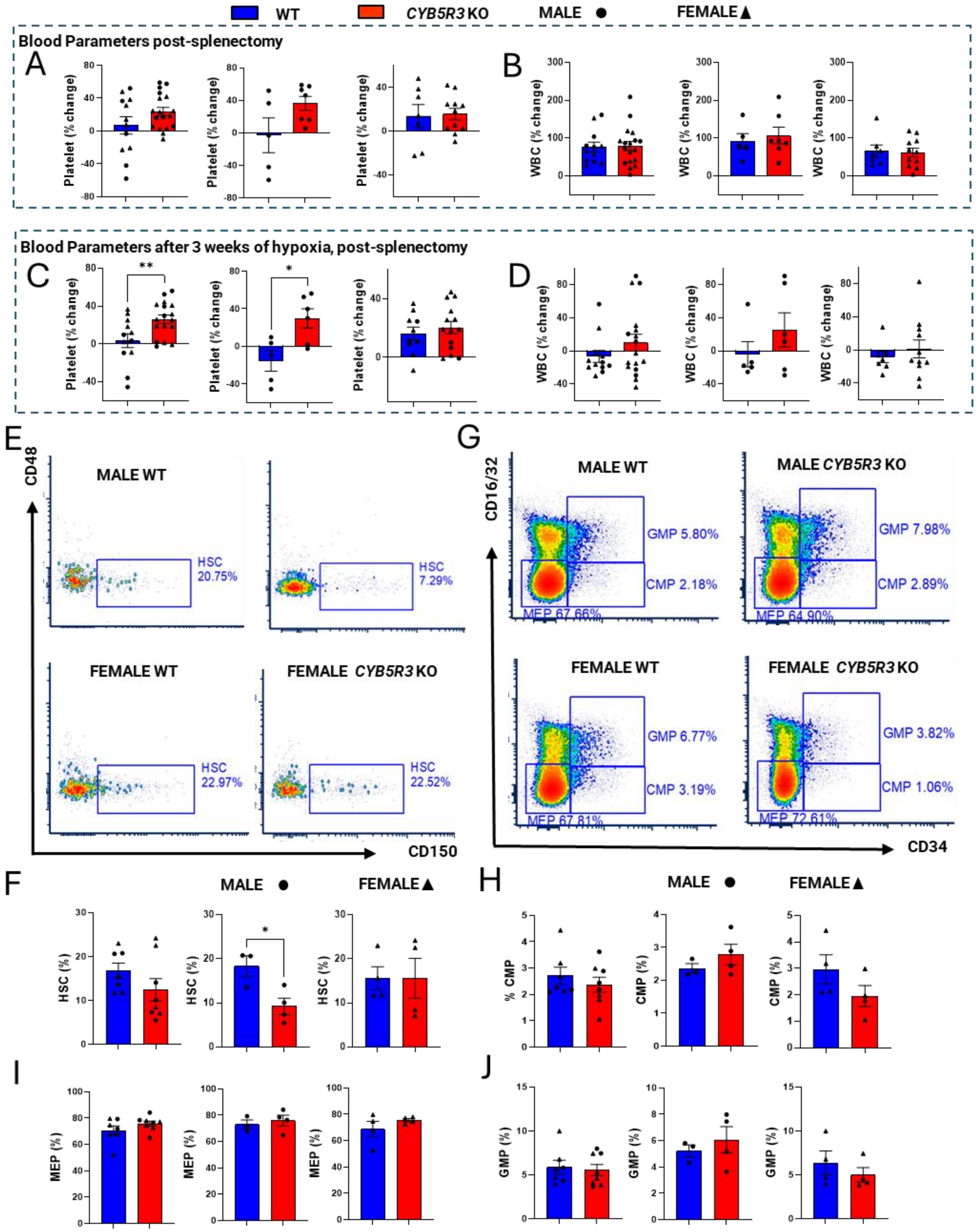

**Splenectomy induces a significant drop in erythropoiesis in CYB5R3 KO mice, which further impacts erythropoietic induction by hypoxia, specifically in male mice.**

**A.** Percentage change in platelets post-splenectomy, combined, and males and females separately. **B.** Percentage change in WBCs post-splenectomy, combined, and males and females separately. **C.** Percentage change in platelets after 3 weeks of hypoxia, post-splenectomy, combined, and males and females separately. **D.** Percentage change in WBCs after 3 weeks of hypoxia, post-splenectomy, combined, and males and females separately. **E.** Flow cytometry plots showing the percentage of HSC population at baseline in the bone marrow after 3 weeks of hypoxia, post-splenectomy. **F.** Quantification of HSCs in the bone marrow, combined, and males and females separately. **G.** Flow cytometry plots showing the percentage of different types of hematopoietic progenitors in the bone marrow after 3 weeks of hypoxia, post-splenectomy. **H-J.** Quantification of different types of hematopoietic progenitors in the bone marrow, combined, and males and females separately. Flow cytometry was performed using a BD Biosciences LSR II. Data were analyzed using FCS Express 7 Research Edition (De Novo Software). GraphPad Prism version 10 was used for statistical analyses. Data are stated as mean ± standard error of the mean unless indicated otherwise. Parametric and non-parametric data with 1 variable were analyzed using Student's t-test and Mann-Whitney test, respectively. F-test was used to compare variances. Grouped data with 2 variables were analyzed using 2-way ANOVA. * indicates $p<0.05$ and was considered significant. CYB5R3: Cytochrome B5 reductase 3; WT: Wild-type; KO: Knockout; WBC: white blood cells; HSC: hematopoietic stem cells; CMP: Common Myeloid Progenitors; MEP: Megakaryocyte Erythroid Progenitors; GMP: Granulocyte Monocyte Progenitors

**Supplemental Figure 6**

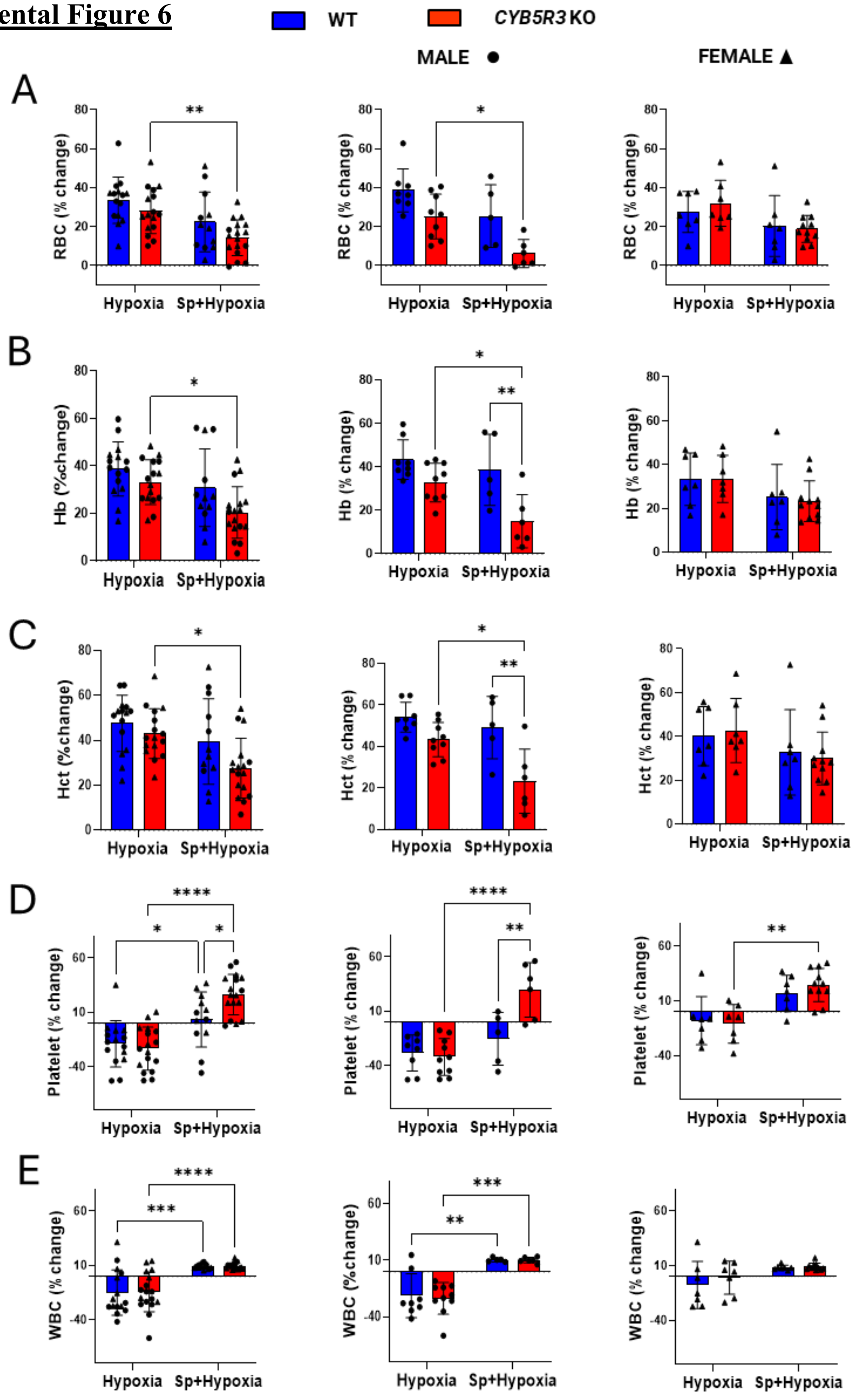


**Splenectomy exacerbates the CYB5R3 knockout effect on stress erythropoiesis, more severely in male mice.**

**A.** Percentage change in RBCs comparing hypoxia vs. hypoxia after splenectomy, combined, and males and females separately. **B.** Percentage change in Hb comparing hypoxia vs. hypoxia after splenectomy, combined, and males and females separately. **C.** Percentage change in Hct comparing hypoxia vs. hypoxia after splenectomy, combined, and males and females separately. **D.** Percentage change in platelets comparing hypoxia vs. hypoxia after splenectomy, combined, and males and females separately. **E.** Percentage change in WBCs comparing hypoxia vs. hypoxia after splenectomy, combined, and males and females separately. GraphPad Prism version 10 was used for statistical analyses. Data are stated as mean ± standard error of the mean unless indicated otherwise. Parametric and non-parametric data with 1 variable were analyzed using Student's t-test and Mann-Whitney test, respectively. F-test was used to compare variances. Grouped data with 2 variables were analyzed using 2-way ANOVA. * indicates $p<0.05$ and was considered significant. CYB5R3: Cytochrome B5 reductase 3; WT: Wild-type; KO: Knockout; Sp: Splenectomy; RBC: red blood cells, erythrocytes; WBC: white blood cells; MEP: Megakaryocyte Erythroid Progenitors; Hb: hemoglobin; Hct: hematocrit

### Supplemental Figure 7

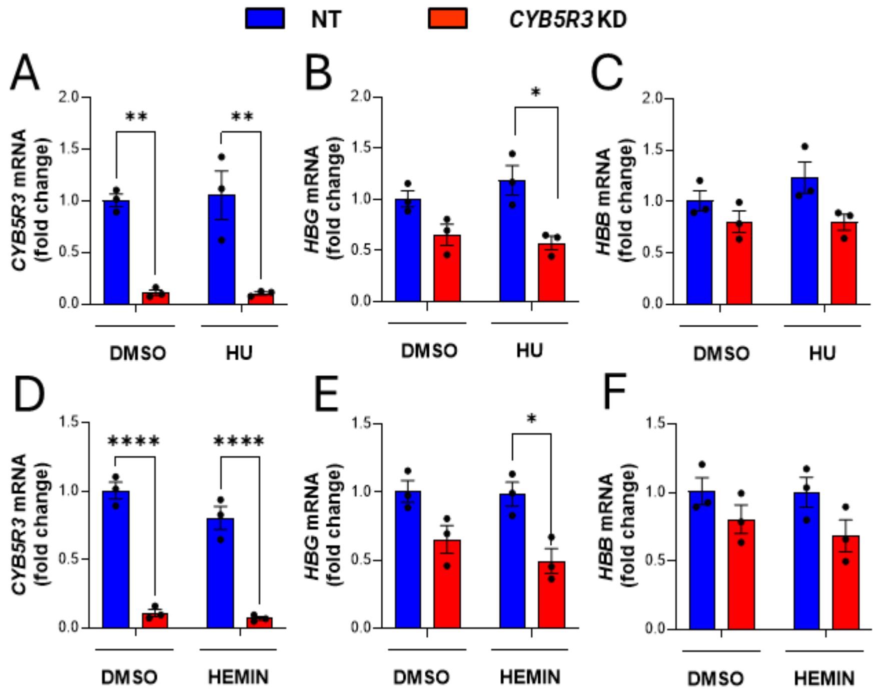


**Knocking down CYB5R3 in K562 cells lowers the globin level, which cannot be rescued by hydroxyurea due to heme deficiency.**

**A-C** Quantification of CYB5R3 and globin mRNA levels after 6 hours of treatment with vehicle or HU. **D-F** Quantification of CYB5R3 and globin mRNA levels after 6 hours of treatment with vehicle or hemin. GraphPad Prism version 10 was used for statistical analyses. Data are stated as mean ± standard error of the mean unless indicated otherwise. Parametric and non-parametric data with 1 variable were analyzed using Student's t-test and Mann-Whitney test, respectively. F-test was used to compare variances. Grouped data with 2 variables were analyzed using 2-way ANOVA. * indicates $p<0.05$ and was considered significant. CYB5R3: Cytochrome B5 reductase 3, NT: Non-targeting, KD: Knockdown; HBG: Gamma globin; HBB: Beta Globin

**Supplemental Figure 8**

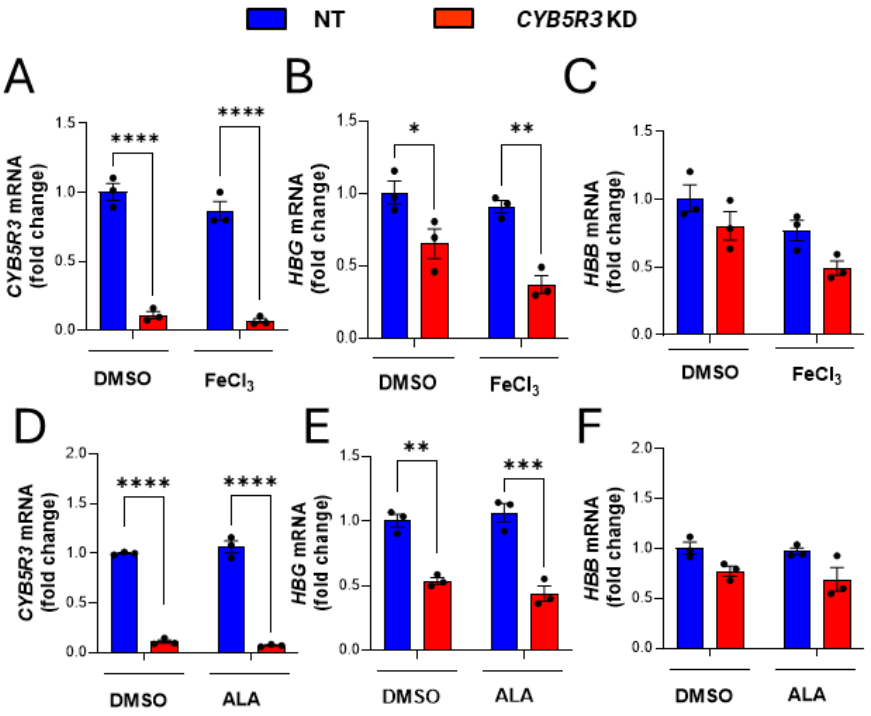


**Globin depletion due to heme deficiency cannot be rescued entirely by iron or a porphyrin intermediate.**

**A-C** Quantification of CYB5R3 and globin mRNA levels after 6 hours of treatment with vehicle or $FeCl_3$. **D-F** Quantification of CYB5R3 and globin mRNA levels after 6 hours of treatment with vehicle or ALA. GraphPad Prism version 10 was used for statistical analyses. Data are stated as mean ± standard error of the mean unless indicated otherwise. Parametric and non-parametric data with 1 variable were analyzed using Student's t-test and Mann-Whitney test, respectively. F-test was used to compare variances. Grouped data with 2 variables were analyzed using 2-way ANOVA. * indicates $p<0.05$ and was considered significant. CYB5R3: Cytochrome B5 reductase 3, NT: Non-targeting, KD: Knockdown; HBG: Gamma globin; HBB: Beta Globin; $FeCl_3$: ferric chloride; ALA: 5-Aminolevulinic acid

**Supplemental Table 1: List of flow antibodies**

| **Antibody-Fluorochrome** | **Catalog no.** |
|---|---|
| **CD44 PE-Cy7** | Biolegend 103030 |
| **CD71 PE** | Biolegend 334106 |
| **TER-119 FITC** | Biolegend 116206 |
| **Streptavidin PerCP-Cy5.5** | Biolegend 405214 |
| **Sca-1 PE-Cy7** | Biolegend 122514 |
| **c-kit APC-Cy7** | Biolegend 105826 |
| **CD150 APC** | Biolegend 115910 |
| **CD48 FITC** | BD Bio 557484 |
| **CD16/32 PE** | LifeTech 12-0161-81 |
| **CD34 FITC** | BD Bio 560238 |
| **Mouse hematopoietic lineage panel Biotin** | LifeTech 88-7774-75 |

**Supplemental Table 2: List of primers**

| Primer | Sequence |
|---|---|
| **Hu ALAS2 Fwd** | CCAAACTGTTCTCAAATCCACC |
| **Hu ALAS2 Rev** | AATCTTGCTCTTCCCATCCTG |
| **Hu ALAD Fwd** | AACCTCATCTACCCCATCTTTG |
| **Hu ALAD Rev** | AAGATCAAGACACAGCGTAGG |
| **Hu PBGD Fwd** | ATTGAAAGCCTCGTACCCTG |
| **Hu PBGD Rev** | TTCTCCAGGGCATGTTCAAG |
| **Hu UROS Fwd** | GAGAAACCTGTGGAAATGCAG |
| **Hu UROS Rev** | ATCCCTTTGTCCTTGAGCG |
| **Hu UROD Fwd** | GACCCCTGTGCCTTGTATG |
| **Hu UROD Rev** | TGTGGTCCAAAGTCATCCAG |
| **Hu CPOX Fwd** | CCAAGAATCCTCATGCTCCTAC |
| **Hu CPOX Rev** | GTGAAAATGGACAGCGTCTTC |
| **Hu PPOX Fwd** | GCAAACCCATCGTTCCATATTAC |
| **Hu PPOX Rev** | CAACATCTCTAGACCTCCACG |
| **Hu HBG Fwd** | AATGTGGAAGATGCTGGAGGAGAA |
| **Hu HBG Rev** | CTTCTTGCCATGTGCCTTGACTT |
| **Hu HBB Fwd** | TCTGTCCACTCCTGATGCTGTTA |
| **Hu HBB Rev** | ATTAGCCACACCAGCCACCACT |
| **Hu HBA Fwd** | TGGACCCGGTCAACTTCAA |
| **Hu HBA Rev** | GAGGCTCCAGCTTAACGGTATT |
| **Hu CyB5R3 Fwd** | ATGGGGGCCCAGCTCAG |
| **Hu CyB5R3 Rev** | AGCGCTGGAACAGCTTCAT |
| **GAPDH Fwd** | ATGACATCAAGAAGGTGGTG |
| **GAPDH Rev** | CATACCAGGAAAATGAGCTTG |
| **Hu GATA1 Fwd** | CTGTCCCCAATAGTGCTTATGG |
| **Hu GATA1 Rev** | GAATAGGCTGCTGAATTGAGGG |

Hu: Human, Fwd: Forward primer, Rev: Reverse primer

**<u>Supplemental Table 3: List of primary antibodies</u>**

| **Antibody** | **Catalog no.** |
|---|---|
| **CYB5R3** | 10894-1-AP |
| **HBA** | sc-514378 |
| **HBB** | sc-21757 |
| **HBB** | 84934S |
| **HBG** | sc-21756 |
| **HBG** | NB110-41084 |
| **Beta (β) actin** | sc-47778 |

### **Extended Methodology**

*Animal experiment at baseline*

A complete blood count (CBC) measurement was conducted using a Heska Analyzer on 12-16 weeks old male and female age-matched WT and *CYB5R3* KO. The bleeding was done retro-orbitally. The 2 groups of mice were then euthanized, and 4 leg bones (2 tibiae and 2 femora) and spleen were harvested from each mouse. The spleen weight was measured and normalized to tibia length. Fresh bone marrow cells were isolated from the bones and used to run a series of flow cytometry experiments to quantify the population of hematopoietic stem cells, erythroid-directed progenitors, and erythroblasts.

*Animal experiment using hypoxia*

12-16 weeks old male and female WT and *CYB5R3* KO mice were used to obtain a baseline CBC using Heska before putting the mice in hypoxia chambers with 10% Oxygen and 90% Nitrogen supply. The mouse CBC was measured weekly for 3 weeks. After a 3-week exposure to hypoxia, the mice were euthanized. 4 leg bones (2 tibiae and 2 femora) and spleen from each mouse were collected. The spleen weight was measured and normalized to tibia length. Afterward, fresh bone marrow cells were collected to perform flow cytometry assays.

*Animal experiment with splenectomy followed by hypoxia*

12-16 weeks old, male and female, WT and *CYB5R3* KO mice underwent a baseline CBC measurement using Heska. Afterward, splenectomy was performed as follows: The mouse was anesthetized (100 mg/kg ketamine and 10 mg/kg xylazine intraperitoneally) and the body temperature was maintained at 37°C using a heating pad. The abdomen was shaved and sterilized, and the abdominal cavity was cut open with a lateral incision on the left side, on a disinfected platform. The splenic vascular pedicles were ligated using 6-0 silk suture, followed by cutting the spleen free from the surrounding tissues before the abdominal incision was closed using a 6-0 silk suture. The mice were provided with 0.1% neomycin water for 2 weeks afterward to prevent any surgery-induced infection. 2 months post-splenectomy, another set of CBCs was done, and the mice were exposed to 3 weeks of hypoxia (10% Oxygen and 90% Nitrogen). The mouse CBC was measured weekly for 3 weeks. The mice were eventually euthanized, and 4 leg bones (2 tibiae and 2 femora) were collected from each mouse. Fresh bone marrow cells were collected to perform flow cytometry assays.

*Electroporation to knock out (KO) CYB5R3 gene in CD34+ HSCs*

On day 3, approximately 250,000 CD34+ HSCs were electroporated using Amaxa Human Stem Cell Nucleofector Kit 2 (VPH-5022, Lonza) in an Amaxa Biosystems Nucleofector II. At room temperature, 20 pmol spCas9 (Synthego) and 100 pmol sgRNA (Synthego) were mixed and incubated for 10 minutes before adding the mixture to the cells for electroporation. The following sgRNAs were used:

NT (non-targeting control): GCAGCTCGACCTCAAGCCGT

KO (knockout): GCTTCATGAGCAGACTGTAC

*Lentiviral transduction for CYB5R3 gene knockdown (KD) in K562 cells*

Lentivirus was generated using the Life Technologies ViraPower Lentiviral System. Briefly, HEK293 cells were transfected with a combination of packaging plasmids and either pLKO.1-puro NT shRNA control plasmid or MISSION pLKO.1-puro containing shRNA targeting the human *CYB5R3* gene (Sigma Millipore SHCLNG TRCN0000236407). Transfected cells were grown for 3 days to allow virus production. The lentivirus was harvested by collecting and concentrating the growth media via centrifugation in concentrator columns (Millipore cat# UFC910008). The NT or KD shRNA sequences used are as follows:

NT: CGCGATAGCGCTAATAATTT

KD: CTGACGCTGCATGAGACATTG

K562 cells were maintained at 0.5-1.0$^{6}$/ml in a 6-well plate. Lentivirus was added to the cells using transduction media made up of growth media supplemented with 10µg/ml polybrene (Millipore TR-1003-G) for a 48-hour incubation. The cells were transferred to a 10 cm dish afterward, and 1µg/ml puromycin was added for the selection. The selection phase continued for a week, splitting the cells as necessary. The cells were either stored or cultured without the selection antibiotic afterward.

*Flow cytometry*

CD34+ HSCs were collected on day 18 and incubated with CD44 antibody for 1 hour in the dark at 4 °C, to analyze the mature erythrocyte population. For analyzing erythroblast stages, mouse bone marrow cells were stained with CD71 and TER-119 and incubated for 1 hour in the dark at 4°C. In both cases, the cells were washed before being taken to run flow cytometry assays. Alternatively, the erythrocytes were removed from the mouse bone marrow cells using a lab-made hemolysis buffer (155mM ammonium chloride, 10mM potassium hydrogen carbonate, 0.1mM EDTA). 2% FBS in PBS was used as a staining buffer to block any Fc receptors. Cells were first stained for 1 hour, as before, with Biotin-conjugated Mouse hematopoietic lineage cocktail to help better selection of lineage-negative cells. The cells were then washed and stained with antibodies against Streptavidin, Sca1, c-kit, CD48, CD150, CD16/32, and CD34. After the incubation period with the antibodies, the cells were washed again. Flow cytometry was performed using a BD Biosciences LSR II. Data were analyzed using FCS Express 7 Research Edition (De Novo Software). Gating was done in a way to select only the live singlets for analysis. **Supp. Table 1** shows the list of flow antibodies used.

*Quantification of gene expression*

K562 cells were collected after 6 hours of each treatment to extract total RNA using an RNA Miniprep Plus kit (Zymo, R2072), according to the manufacturer's instructions. SuperScript™ IV VILO™ Master Mix (Thermo Fisher Scientific, 11756050) was used for reverse transcription to make cDNA. Power SYBR™ Green PCR Master Mix (Thermo Fisher Scientific, 4367659) and QuantStudioTM 5 real-time PCR machine (Thermo Fisher Scientific) were used for conducting quantitative PCR using cDNA. The mRNA levels were measured using the delta-delta Ct method. Approximately 2 μg RNA was used for bulk RNA sequencing performed by Novogene, as per company recommendations, to identify differentially expressed genes and pathways between the experimental groups. **Supp. Table 2** shows the list of primers used.

*RNA sequencing*

K562 cells were collected after 6 hours treatment with DMSO to extract total RNA using an RNA Miniprep Plus kit (Zymo, R2072), according to the manufacturer's instructions. Total RNA was then sent to Novogene for unstranded library preparation and bulk RNA sequencing (150 bp paired) on an Illumina platform. Raw FASTQ files were filtered to remove low-quality reads using fastq and aligned to the human genome using Hisat2 v2.0.5. Mapped reads are then counted using featureCounts v1.5.0-p3 on the gene level. The downstream bioinformatic analysis was performed using R 4.4.1. From the gene count data, low-count genes with less than 1 count per million reads (CPM) were removed. The filtered gene counts were used by DESeq2 1.44.0 for differential gene expression analysis with the Wald test. Genes with adjusted p-values (false discovery rate) $< 0.05$ were considered significant. We then performed over-representation pathway analysis with ClusterProfiler 4.12.6 using differentially expressed genes and the Gene Ontology database.

*Protein analysis and quantification*

Cultured cells were lysed by sonication in 1X RIPA buffer supplemented with protease (Millipore Sigma, P8340) and phosphatase inhibitors (Millipore Sigma, P5726). Lysates were quantified using BCA assay (Thermo Fisher, 23225) and prepared in 1X Laemmli buffer by boiling at 100°C for 5 minutes. 12-15µg protein was loaded in each well of 4%–12% Bis-Tris SDS Gel (Life Tech, NP0321BOX) for electrophoresis at 180V for 65 minutes using MES Running Buffer. The proteins were transferred at 100V for 1 hour in Tris-Glycine buffer to nitrocellulose membranes. Membranes were blocked in 1% BSA in PBS for 1 hour and then incubated in primary antibody diluted in 1% BSA in PBST (0.1% Tween PBS pH 7.4) overnight in a cold room. LICOR secondary antibodies were used to blot the membranes for 1 hour at room temperature, followed by signal detection using the LI-COR Odyssey imaging system. **Supp. Table 3** shows the list of primary antibodies used.

*Sample Preparation and HPLC-MS/MS Analysis*

Cell pellets were utilized to quantify the levels of protoporphyrinogen IX and heme. To prepare the samples, cell pellets were spiked with 33.4 pmol of deuteroporphyrin IX 2-vinyl, 4-hydroxymethyl as an internal standard. Deproteinization was conducted by adding 100µl of 80% cold methanol and centrifugation at 10,000 rpm at 4°C for 10 minutes. The resulting supernatant was transferred to a new vial for subsequent HPLC-MS/MS analysis. Solvents used for extractions and mass spectrometric analyses were HPLC grade or better from Fisher Scientific (Fairlawn, NJ).

Protoporphyrinogen IX and heme were analyzed by HPLC-ESI-MS/MS using a gradient solvent system consisting of water containing 0.1% formic acid (solvent A) and acetonitrile containing 0.1% formic acid (solvent B). A reverse phase HPLC column (2 × 100 mm x 5 µm Luna C18(2) column; Phenomenex) was used with a flow rate of 0.65 ml/min. Samples were loaded onto the column at 10% B, held for 0.3 min, and eluted with a linear increase in solvent B from 10 to 100% over 8.7 min. Analyte quantification was carried out in multiple reaction monitoring (MRM) mode using a QTrap 6500+ triple quadrupole mass spectrometer (Sciex, San Jose, CA) equipped with an electrospray ionization source operating in the positive-ion mode. The instrument parameters were set as follows: collision gas at 5 units, curtain gas at 40 units, ion source gas number 1 at 55 units, and number 2 at 60 units. The ion spray voltage was set at 5500 V, and the temperature was maintained at 600 °C. The declustering potential, entrance potential, collision energy, and collision exit potential were set to 85 eV, 10, 55, and 10, respectively. MRM transitions utilized were as follows: 563.3/504.4 for protoporphyrin IX, 616.1/557.1 for heme, and 567.3/508.3 for deuteroporphyrin IX 2-vinyl, 4-hydroxymethyl.
